# Pathogenic diversification of the gut commensal *Providencia alcalifaciens* via acquisition of a second type III secretion system

**DOI:** 10.1101/2024.06.07.595826

**Authors:** Jessica A. Klein, Alexander V. Predeus, Aimee R. Greissl, Mattie M. Clark-Herrera, Eddy Cruz, Jennifer A. Cundiff, Amanda L. Haeberle, Maya Howell, Aaditi Lele, Donna J. Robinson, Trina L. Westerman, Marie Wrande, Sarah J. Wright, Nicole M. Green, Bruce A. Vallance, Michael McClelland, Andres Mejia, Alan G. Goodman, Johanna R. Elfenbein, Leigh A. Knodler

**Affiliations:** Paul G. Allen School for Global Health, College of Veterinary Medicine, Washington State University, Pullman, WA, USA; Wellcome Sanger Institute, Hinxton, Cambridgeshire, United Kingdom; Department of Pathobiological Sciences, School of Veterinary Medicine, University of Wisconsin-Madison, Madison, WI, USA; Department of Clinical Sciences, College of Veterinary Medicine, North Carolina State University, Raleigh, NC, USA; Public Health Laboratory, Los Angeles County Department of Public Health, CA; Division of Gastroenterology, Department of Pediatrics, BC Children’s Hospital and the University of British Columbia, Vancouver, BC, Canada; Department of Microbiology and Molecular Genetics, University of California, Irvine, CA, USA; Comparative Pathology Laboratory, Research Animal Resources and Compliance, University of Wisconsin-Madison, Madison, WI, USA; School of Molecular Biosciences, College of Veterinary Medicine, Washington State University, Pullman, WA, USA; Department of Microbiology and Molecular Genetics, Robert Larner College of Medicine at The University of Vermont, Burlington, VT, USA

**Author notes:** These authors contributed equally (author order was determined alphabetically by surname) (Mattie M. Clark-Herrera, Eddy Cruz, Jennifer A. Cundiff, Amanda L. Haeberle, Maya Howell, Aaditi Lele, Donna J. Robinson, Trina L. Westerman, Marie Wrande, Sarah J. Wright). Complex *in vitro* Systems, Safety Assessment, Genentech Inc., South San Francisco, CA, USA. Department of Medical Biochemistry and Microbiology, Uppsala University, Uppsala, Sweden (Marie Wrande).

## Abstract

*Providencia alcalifaciens* is a Gram-negative bacterium found in a wide variety of water and land environments and organisms. It has been isolated as part of the gut microbiome of animals and insects, as well as from stool samples of patients with diarrhea. Specific *P. alcalifaciens* strains encode gene homologs of virulence factors found in other pathogenic members of the same Enterobacterales order, such as *Salmonella enterica* serovar Typhimurium and *Shigella flexneri.* Whether these genes are also pathogenic determinants in *P. alcalifaciens* is not known. Here we have used *P. alcalifaciens* 205/92, a clinical isolate, with *in vitro* and *in vivo* infection models to investigate *P. alcalifaciens*-host interactions at the cellular level. Our particular focus was the role of two type III secretion systems (T3SS) belonging to the Inv-Mxi/Spa family. T3SS_1b_ is widespread in *Providencia* spp. and encoded on the chromosome. T3SS_1a_ is encoded on a large plasmid that is present in a subset of *P. alcalifaciens* strains, which are primarily isolates from diarrheal patients. Using a combination of electron and fluorescence microscopy and gentamicin protection assays we show that *P. alcalifaciens* 205/92 is internalized into eukaryotic cells, rapidly lyses its internalization vacuole and proliferates in the cytosol. This triggers caspase-4 dependent inflammasome responses in gut epithelial cells. The requirement for the T3SS_1a_ in entry, vacuole lysis and cytosolic proliferation is host-cell type specific, playing a more prominent role in human intestinal epithelial cells as compared to macrophages. In a bovine ligated intestinal loop model, *P. alcalifaciens* colonizes the intestinal mucosa, inducing mild epithelial damage with negligible fluid accumulation. No overt role for T3SS_1a_ or T3SS_1b_ was seen in the calf infection model. However, T3SS_1b_ was required for the rapid killing of *Drosophila melanogaster*. We propose that the acquisition of two T3SS by horizontal gene transfer has allowed *P. alcalifaciens* to diversify its host range, from a highly virulent pathogen of insects to an opportunistic gastrointestinal pathogen of animals.

## Introduction

Despite belonging to a large and clinically significant family of Gram-negative bacteria, *Providencia* species remain among the least studied members of the Enterobacteriaceae. *Providencia* spp. colonize diverse hosts and environments. In addition to being found in soil (1), water (2), sewage (3) and retail meats, fruits and vegetables (4–7), *Providencia* spp. are members of the human gut, oral cavity and sputum microbiomes (8–12). *Providencia* spp. have also been isolated from numerous animals including penguins, turtles, sharks, snakes (13), nematodes (14), and insects such as blow flies, fruit flies, house flies and olive flies (13, 15–18). Notably, some *Providencia* spp. are pathogenic to *Drosophila melanogaster, Ceratitis capitata* (Mediterranean fruit fly) and *Anastrepha ludens* (Mexican fruit fly), with the most highly virulent species being *Providencia alcalifaciens* and *Providencia sneebia* in *D. melanogaster* (13) and *P. alcalfaciens* and *Providencia rustigianii* in *A. ludens* (19).

Considered as opportunistic bacterial pathogens of humans, *P. alcalifaciens*, *Providencia rettgeri*, and *Providencia stuartii* are the most common clinical isolates (20, 21) and cause a spectrum of nosocomial and environmentally acquired diseases, including urinary tract, wound and ocular infections, as well as diarrhea, meningitis and sepsis. *Providencia* spp. are typically resistant to the penicillins, first-generation cephalosporins, aminoglycosides, tetracyclines and polymyxins (22, 23). Their increasing antimicrobial resistance is a major public health concern (24, 25). Several groups have reported *P. alcalifaciens* to be a cause of diarrhea in infants and travelers in developing countries (21, 26–30) and in foodborne-associated outbreaks (31–33). The incidence of *P. alcalifaciens* in diarrheal patients in Thailand (1.9%), Bangladesh (2.1%) and Kenya (3.2%) is on par with *Salmonella* spp. (4, 29, 34). A higher incidence of *P. alcalifaciens* (10-18%) has been reported for persons with traveler’s diarrhea (21, 35). *P. alcalifaciens* has also been associated with diarrhea in dogs and cats (36–39). Previous work has verified the ability of some *P. alcalifaciens* clinical isolates to elicit diarrheal disease in a removable intestinal tie adult rabbit diarrhea (RITARD) infection model (40, 41) and cause fluid accumulation in rabbit ileal loops (31). Furthermore, clinical strains isolated from patients with diarrhea exhibit varying invasive abilities with some *P. alcalifaciens* being highly invasive for human epithelial cell lines e.g. HeLa, HEp-2, Vero and Caco-2, and others being non-invasive (31, 42–48). Despite a strong association with diarrheal illness in humans and animals, a detailed understanding of the pathogenic mechanisms of *P. alcalifaciens* is lacking.

Enteric pathogens such as *Salmonella enterica* serovar Typhimurium (*S*. Typhimurium) and *Shigella flexneri* use type III secretion systems (T3SSs), also known as injectisomes, to deliver “effector” proteins into host cells that modulate the actin cytoskeleton, allowing for their efficient entry into non-phagocytic cells. T3SSs are found in many (but not all) *Providencia* isolates, including *P. alcalifaciens* (49), an indication of the pathogenic potential of members of this genus. We reported earlier that *P. alcalifaciens* 205/92, a clinical isolate, encodes for two T3SSs belonging to the Inv-Mxi/Spa family, which we designated T3SS_1a_ and T3SS_1b_ (50). *Sodalis glossinidius* (51), an insect endosymbiont, and some isolates of *Providencia* spp. are the only other bacteria known to encode for two Inv-Mxi/Spa T3SS. *P. alcalifaciens* T3SS_1a_ is closely related to, and functionally interchangeable with, the invasion-associated T3SS1 from *S.* Typhimurium (50). Structural proteins of T3SS_1b_ share significant amino acid sequence identity to those of the Ysa T3SS from *Yersinia enterocolitica* (50), which is restricted to biotype 1B (52). Neither Ysa nor T3SS_1b_ translocator operons functionally substitute for those of *S.* Typhimurium in driving bacterial entry into non-phagocytic cells, suggesting an evolutionary functional divergence within the Inv-Mxi/Spa family of T3SSs (50).

We hypothesized that T3SSs are virulence determinants in *P. alcalifaciens*. Here we have investigated the role of the two T3SSs in bacterial colonization of mammalian and insect cells, specifically human intestinal epithelial cells (IECs), human macrophages and *D. melanogaster* macrophage-like cells. We report that *P. alcalifaciens* 205/92 enters eukaryotic cells, rapidly lyses its internalization vacuole, then replicates within the cytosol. We further show that T3SS_1a_, which is encoded on a 128 kb plasmid, is necessary for efficient bacterial entry, nascent vacuole lysis, and intracellular replication in human IECs. In human macrophages, a T3SS_1a_ mutant induces less host cell cytotoxicity than wild type bacteria. While a T3SS_1b_ mutant exhibits no colonization defect in mammalian or insect cell lines, it is significantly attenuated for infection of *D. melanogaster*. Therefore, *P. alcalifaciens* 205/92 uses two type III injectisomes to colonize diverse eukaryotic hosts.

## Results

### P. alcalifaciens *205/92 genome*

Seven *Providencia alcalifaciens* genomes have been sequenced as part of the NIH Common Fund Human Microbiome Project (53) and deposited in GenBank, including *P. alcalifaciens* 205/92 (https://www.ncbi.nlm.nih.gov/datasets/genome/GCF_000527335.1/). This strain was originally isolated from a young Bangladeshi boy with diarrhea (42, 45). The contig-level genome assembly includes 88 contigs. We generated a complete *P. alcalifaciens* 205/92 genome using a combination of Oxford Nanopore Technologies (ONT) long read and Illumina short read sequencing. Assembly of the highest quality 100x filtered and trimmed Nanopore reads (see Methods section) using Trycycler generated three circular replicons: a 4,094,134 bp chromosome, and two plasmids, 127,796 bp (p128kb) and 40,541 bp (p40kb) in size. Due to size selection, smaller plasmids are often under-represented in long read-only assemblies. To account for this, we performed an independent hybrid assembly of the Nanopore reads with bbduk-trimmed Illumina reads which allowed us to recover a third plasmid, 3,997 bp in size (p4kb). p128kb and p40kb are single-copy plasmids, whereas p4kb is multi-copy. Polishing the resulting combined assembly with Polypolish using the bbduk-trimmed Illumina reads corrected three errors in the chromosome, and two in the 128kb plasmid. The GenBank Accession number is GCA_038449115.1 (https://www.ncbi.nlm.nih.gov/datasets/genome/GCA_038449115.1/).

The *P. alcalifaciens* 205/92 complete genome has a G+C content of 41.8%, in line with the average G+C content of *Providencia* spp. genomes (54). Automated annotation of the assembled genome using Bakta identified 4,007 protein coding genes (chromosome: 3,842 protein coding genes; p128kb: 113 protein coding genes; p41kb: 46 protein coding genes; p4kb: 6 protein coding genes), as well as 7 ribosomal operons (22 rRNA genes overall), and 80 tRNA genes. Using the PHASTER prophage-prediction web tool followed by manual curation, we identified nine putative prophage regions (Figure 1A). Genomic islands encoding flagella, a type VI secretion system (T6SS) and a type III secretion system, T3SS_1b_, were present on the chromosome (Figure 1A). Genomic islands associated with a type IV secretion system (T4SS) and a second type III secretion system, T3SS_1a_ (Figure 1B), were found on p40kb and p128kb, respectively.

**Figure 1.**
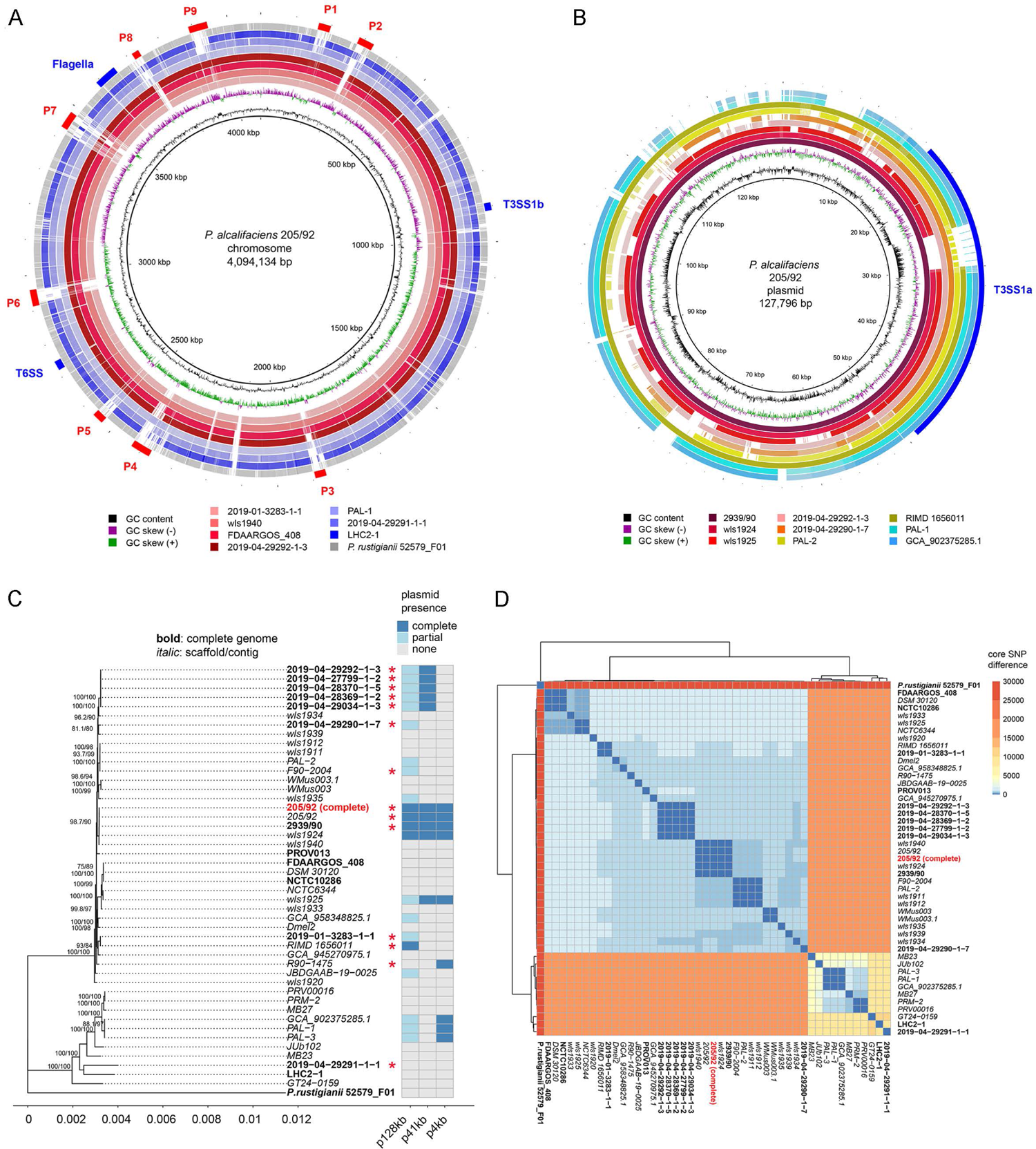
Providencia alcalifaciens 205/92 genome. (A) BRIG diagram of chromosome alignment of seven *P. alcalifaciens strains* (2019-01-3283-1-1, wls1940, FDAARGOS_408, 2019-04-29292-1-3, PAL-1, 2019-04-29291-1-1, LHC2-1) and *P. rustigianii* strain 52579_F01. Complete genome of strain 205/92 presented in this work is used as a reference. Shades of red indicate the phylogenetic branch that includes 205/92; shades of blue indicate the second major branch. Exact coordinates of putative prophages P1-P9, flagella, and secretion systems are provided in Table S1. (B) BRIG diagram of p128kb alignment from nine *P. alcalifaciens* strains (2939/90, wls1924, wls1925, 2019-04-29292-1-3, 2019-04-29290-1-7, PAL-2, RIMD 1656011, PAL-1, and GCA_902375285.1). p128kb of the 205/92 strain presented in this work is used as a reference. Shades of blue indicate strains from the second phylogenetic branch that does not include 205/92. Exact coordinates of T3SS_1a_ are provided in Table S1. (C) Whole genome phylogenetic tree generated with IQTree from the core genome alignment of 46 *P. alcalifaciens* strains and *P. rustigianii* strain 52579_F01 used as an outgroup. Bold font indicates complete genome assemblies; italic font indicates genomes assembled into multiple contigs or scaffolds. The heatmap indicates presence (complete or partial) or absence of the three plasmids present in the *P. alcalifaciens* 205/92 genome. Plasmid is classified as complete if read coverage is >80%; as partial if read coverage is 40-80%; and absent if read coverage is <40%. Red asterisks denote isolates collected from human or canine patients with diarrhea (see Table S1 for detailed information on strains). (D) A heatmap of pairwise core SNP distances between the 45 *P. alcalifaciens* strains and *P. rustigianii* strain 52579_F01. The total number of identified high-quality (AGCT-only) variable positions is 44,562.

We compared the *P. alcalifaciens* 205/92 genome with other *P. alcalifaciens* genomes (Figure 1). Core genome alignment identified 44,562 high quality (AGCT-only) variant positions alongside 3,937,741 constant sites. Phylogenetic analysis of the resulting whole-genome alignment using IQTree (www.iqtree.org/doc/Substitution-Models) identified an TVM+F+I+R2 model as best fitting the data according to the Bayesian Information Criterion. *P. rustigianii* strain 52579_F01 was used as an outgroup. Phylogenetic analysis of the strains (Figure 1C) generally confirmed the conclusions made previously based on the protein-coding based phylogeny (49): strains designated as *P. alcalifaciens* could be divided into two highly distinct clusters (Figure 1C). Cluster 1, which includes strain 205/92 and a closely related strain 2939/90, also isolated in Bangladesh from a child with diarrhea (40), has much closer and fewer differences between the sequenced genomes, and far more isolated strains than cluster 2, which could be the result of a fast and recent spread of Cluster 1. We checked for the presence of T3SS_1a_ and T3SS_1b_ in different *P. alcalifaciens* strains that were chosen according to phylogeny from the two major clusters. The chromosomal-encoded T3SS_1b_ was fully present in all the profiled strains (Figure 1A). For the plasmid-encoded T3SS_1a_, our analysis was limited to those strains with the largest plasmid (Figures 1B, 1C). Cluster 1 is characterized by a much higher presence of the two large plasmids identified in the 205/92 strain. Indeed, out of 34 isolates that belong to cluster 1, 4 carried a full and 13 carried a substantial partial representation of the 128 kb plasmid. Notably, there was a strong correlation between isolates from diarrheal patients (humans or dogs) and the presence of p128kb (Figure 1C). These isolates were sourced from diverse geographical locations (Table S1). In contrast, only 3 out of 11 isolates from cluster 2 carried a partial p128kb. We were surprised to find that, despite the considerable differences in the overall genetic content of this plasmid between strains, coding potential for all T3SS structural and regulatory proteins remained intact in all the strains (Figures 1B, S1). Interestingly, there was considerable heterogeneity between strains in the protein-coding sequence length for a predicted type III effector, SipA. In *P. alcalifaciens* 205/92, SipA is a >2,300 AA protein (Figure 1B). Nine out of 34 isolates from cluster 1 and none from cluster 2 carried the 41kb plasmid. The small p4kb plasmid had a more uniform distribution: 5/34 for cluster 1, and 3/11 for cluster 2 (Figure 1C).

Core SNP differences (Figure 1D) also confirm the overall picture described in Yuan et al. (2020). A mean distance between isolates from cluster 1 and cluster 2 was determined to be between 16,000 and 17,000 SNPs per 44,562 high-quality variable positions. Cluster 1 had a mean distance between the two strains of approximately 1,000 SNPs; at the same time, the mean distance between the two strains in cluster 2 was over 5,000 SNPs (Figure 1D). Overall, it can be concluded that the *P. alcalifaciens* species is genomically diverse and consists of two major lineages.

### T3SS_1a_ genes are induced in late-log phase of growth

Given the involvement of secretion systems in the pathogenesis of numerous Gram-negative bacteria, we initially set out to define *in vitro* growth conditions under which *P. alcalifaciens* T3SS genes are transcriptionally active. The promoter regions of *invF, prgH, and sicA*, the first gene in predicted operons from T3SS_1a_ and T3SS_1b_ pathogenicity islands (Figure 2A), were cloned upstream of a promoterless *luxCDABE* in pFU35. The resulting plasmids were electroporated into wild type (WT) *P. alcalifaciens*. We detected robust luminescence for bacteria carrying P*prgH*_1a_-*luxCDABE* and P*invF*_1a_-*luxCDABE* reporters at late log-phase of growth in LB-Miller broth, pH 7.0, at 37°C (Figure 2B, 2C). Transcriptional activity for *sicA*_1a_ was much lower, but still greater than that of the promoter-less vector, pFU35, under these conditions (Figure 2C). By contrast, no luminescence was detected for bacteria carrying T3SS_1b_ gene reporters under any of the *in vitro* growth conditions we tested i.e. shaking in LB-Miller broth, pH 7.0 (Figure 2C) or pH 5.8 at 37°C or 25°C; shaking in M9 minimal media pH 7.0 at 37°C or 25°C; or McCoy’s medium or Schneider’s medium in the absence or presence of 10% heat-inactivated calf serum at 25°C and 37°C (data not shown). Collectively, we conclude that T3SS_1a_- and T3SS_1b_-associated genes are expressed under distinct conditions.

**Figure 2.**
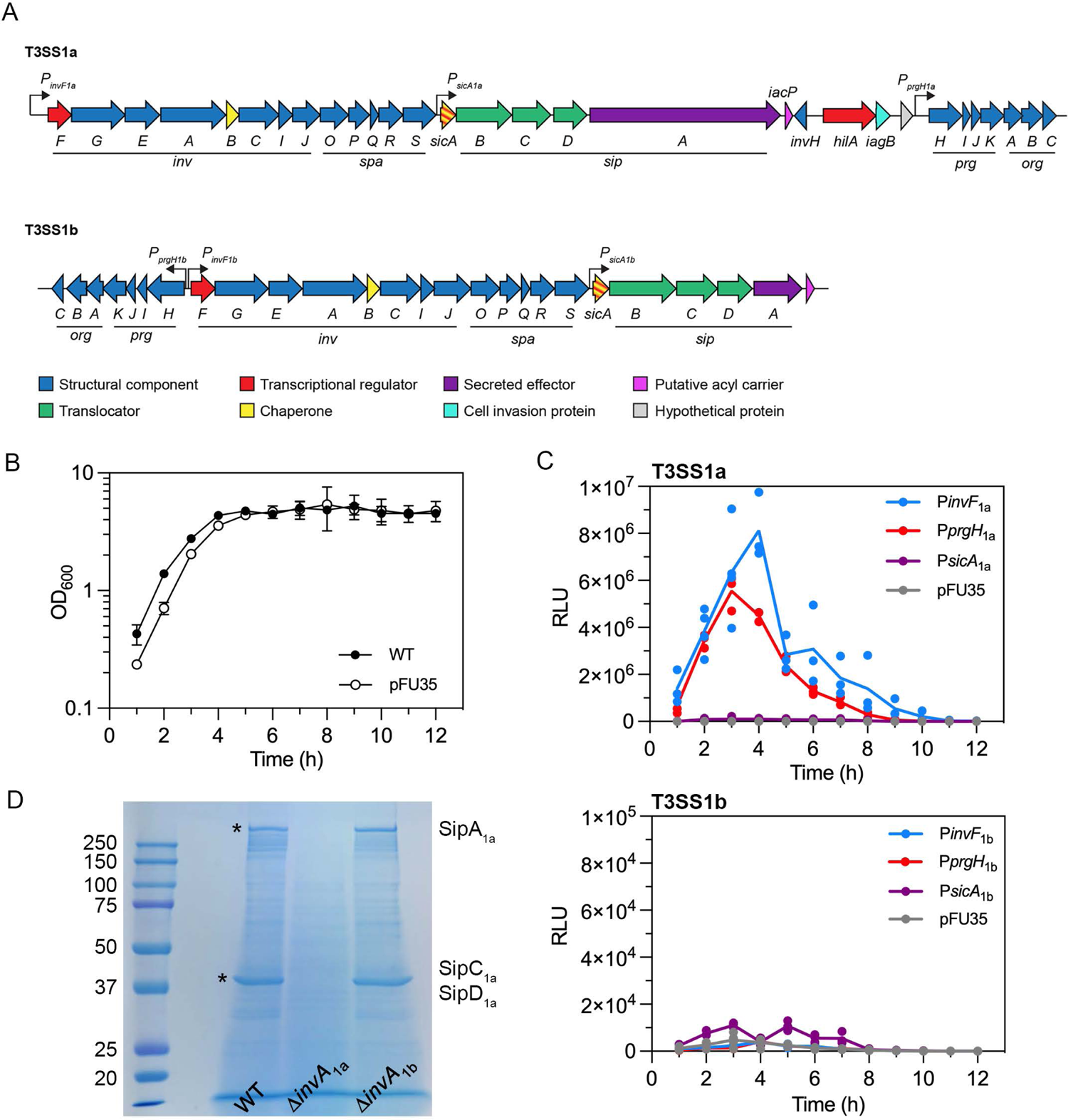
Characterization of T3SS_1a_ and T3SS_1b_ in *P. alcalifaciens* 205/92. (A) Cartoon depiction of the genetic organization of T3SS_1a_ and T3SS_1b_. (B) Growth curve of *P. alcalifaciens* 205/92 WT and WT carrying pFU35 plasmid. Bacterial subcultures were grown at 37°C, shaking at 220 rpm in LB-Miller broth and OD_600_ measured hourly. Mean ± SD of three independent experiments. (C) Bacterial luminescence over a time course of growth. *P. alcalifaciens* 205/92 carrying *luxCDABE* transcriptional reporters for the indicated T3SS_1a_ (upper panel) or T3SS_1b_ (lower panel) gene promoters or the empty plasmid (pFU35) were grown as in (B) and relative light units (RLU) were measured every hour in a microplate reader. Lines indicate the mean of three independent experiments (each symbol represents data from one experiment). (D) Secreted proteins profile. *P. alcalifaciens* WT, Δ*invA*_1a_ and Δ*invA*_1b_ subcultures were grown for 4 h in LB-Miller broth. Supernatants were collected, filtered and precipitated proteins separated by SDS-PAGE and stained with GelCode Blue. Molecular mass markers are shown on the left. The protein bands indicated by asterisks were excised and identified as SipA_1a_ (>250 kDa), SipC_1a_ (∼40 kDa) and SipD_1a_ (∼40 kDa) by mass spectrometry.

*Salmonella* spp. secrete effector proteins into culture media in a type III secretion-dependent manner (55). When *P. alcalifaciens* was grown under *in vitro* conditions that induce T3SS_1a_-associated genes (late-log phase in LB-Miller broth), numerous proteins were secreted into the culture supernatant (Figure 2D). To identify whether protein secretion was dependent on T3SS_1a_ or T3SS_1b_ we compared the protein profiles of culture supernatants from WT, Δ*invA*_1a_ and Δ*invA*_1b_ deletion mutant bacteria. InvA is a highly conserved inner membrane component of the Inv-Mxi/Spa T3SS family. *S*. Typhimurium *invA* mutants are unable to assemble the needle portion of the injectisome and, as a result, are deficient for T3SS-dependent protein secretion (56). By analogy, we presume that *P. alcalifaciens invA* deletion mutants are secretion-incompetent. The secreted protein profile of Δ*invA*_1b_ bacteria was indistinguishable from that of WT bacteria whereas at least two protein bands were absent from supernatants of Δ*invA*_1a_ bacteria, one at >250 kDa and one at ∼40 kDa (Figure 2D). Mass spectrometric analysis identified the proteins as SipA_1a_ (predicted molecular mass of 240 kDa), and a mixture of SipC_1a_ and SipD_1a_ (predicted mass of 43 and 39 kDa, respectively), the corresponding genes of which are encoded on T3SS_1a_ (Figure 2A). Due to the presence of tandem repeat sequences, 205/92 SipA_1a_ is much larger than orthologous proteins from *S*. Typhimurium (SipA), *Shigella flexneri* (IpaA) (∼74kDa) and some other *P. alcalifaciens* strains. Of the 20 strains harboring p128kb, 11 encode for SipA_1a_ of >200 kDa (Figure 1B and Figure S1). In *Salmonella* spp. SipA is a type III effector with actin-binding properties (57), SipC is a type III translocator protein (58) and SipD is the needle tip protein of T3SS1 (58).

*Providencia* spp. have peritrichous flagella and are considered motile. *P. alcalifaciens* 205/92 encodes numerous flagella-associated genes in one large genetic island on the chromosome (Figure 1A, Figure S2A). We constructed *luxCDABE*-based transcriptional reporters to the promoter region of *flhD*, *flgB* and *fliC* and measured bacterial luminescence when WT bacteria carrying these reporters were grown in LB-Miller broth pH 7.0 at 37°C for 12 h. All three genes were transcribed, with a peak of transcription at late log-phase of growth (Figure S2B). Swimming motility of *P. alcalifaciens* was evident on soft agar plates, albeit much less than *S*. Typhimurium WT bacteria but greater than a non-motile *S*. Typhimurium Δ*flgB* mutant (Figure S2C). Overall, we conclude that *P. alcalifaciens* T3SS_1a_ and flagellar genes are induced by aeration and at late log-phase of growth, similar to T3SS1 genes in *S*. Typhimurium (59).

### Internalization of P. alcalifaciens into mammalian and insect cell lines

Some clinical isolates of *P. alcalifaciens*, including 205/92, have been reported to invade HEp-2 cell monolayers (42, 45). To investigate the phenotypic characteristics of the initial interaction of *P. alcalifaciens* 205/92 with mammalian non-phagocytic cells, we used scanning electron microscopy (SEM) and transmission electron microscopy (TEM). *P. alcalifaciens* were grown under conditions that induce the T3SS_1a_ (Figure 2C), added to monolayers (HeLa or HCT116 epithelial cells), centrifuged for 5 min and incubated for a further 15 min (20 min p.i. for SEM) or 55 min (1 h p.i. for TEM) at 37°C, then infected monolayers were processed for microscopy. By SEM, we observed *P. alcalifaciens* attaching to filopodial extensions on the epithelial cell surface (Figure 3A, 3C). Sometimes these membrane protrusions were wrapped around the bacteria (Figure 3A). Similar initial interactions with epithelial cells have been described for *S. flexneri*, *Y. enterocolitica* and *H. pylori* (60–63). Invasion of *P. alcalifaciens* into epithelial cells is associated with actin condensation at the site of bacterial entry (40), and inhibited by cytochalasin D (42), a hallmark of the “trigger” type of cell entry mediated by *S*. Typhimurium and *S. flexneri.* By SEM, we did not observe dramatic plasma membrane ruffles characteristic of bacterial entry via this mechanism, however. TEM analysis suggested *P. alcalifaciens* internalization into non-phagocytic cells was instead via a zipper-like mechanism; upon bacterial adherence, membrane protrusions formed and wrapped around bacteria in tight apposition, eventually engulfing the entire bacterium (Figure 3B, 3D).

**Figure 3.**
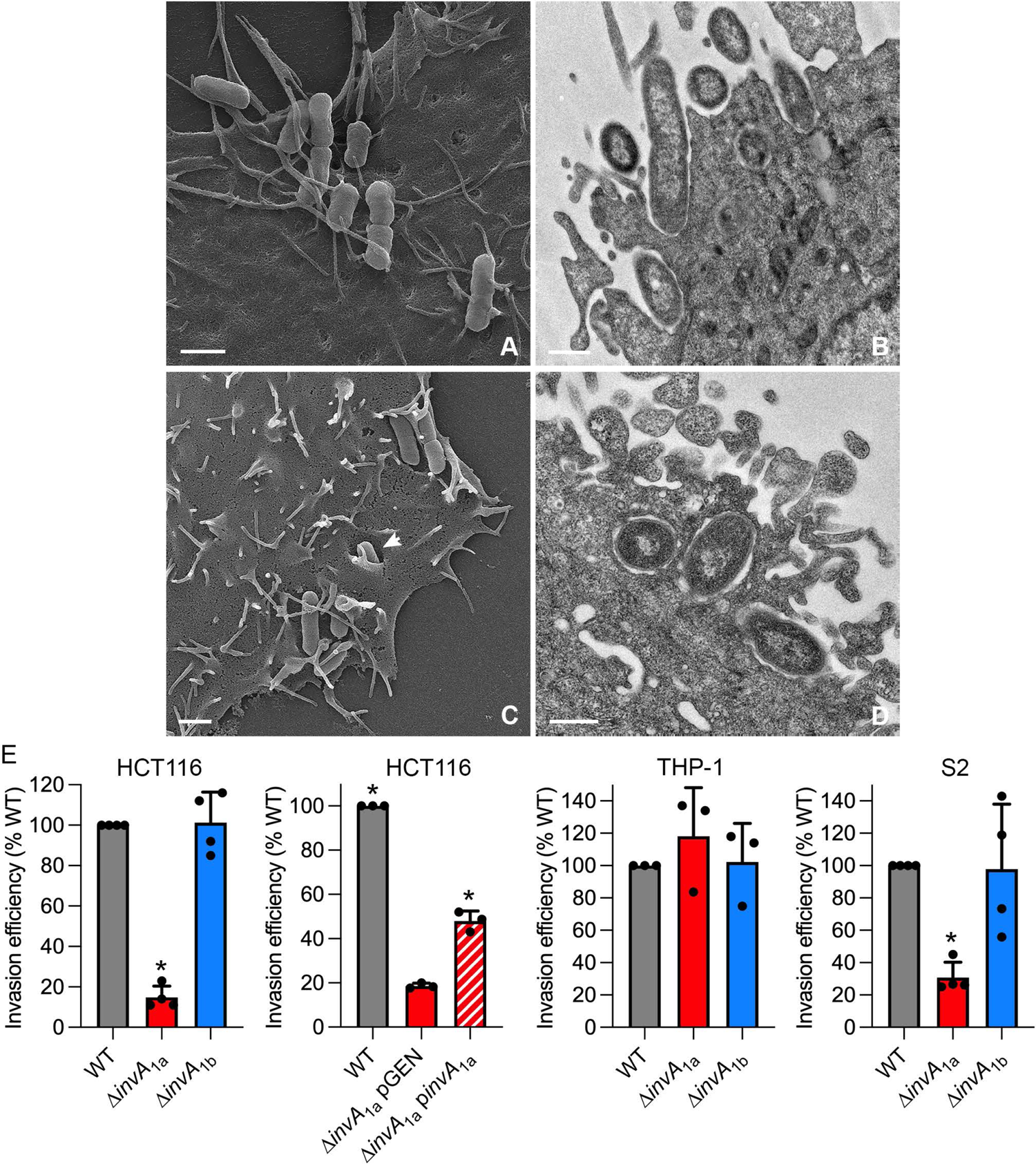
*P. alcalifaciens* adherence to, and invasion of, eukaryotic cells. (A) Scanning electron micrograph (SEM) of *P. alcalfaciens* 205/92 associating with filopodia on the surface of HeLa epithelial cells at 20 min p.i. Scale bar is 1 µm. (B) Transmission electron micrograph (TEM) showing *P. alcalifaciens* entering a HeLa epithelial cell at 1 h p.i. Scale bar is 0.5 µm (C) SEM shows bacteria associating with filopodia on the surface of a HCT116 colonic epithelial cell at 20 min p.i. One bacterium is seen entering via a zipper-like mechanism (indicated by arrowhead). Scale bar is 1 µm. (D) TEM showing *P. alcalfaciens* entering a HCT116 cell at 1 h p.i. Scale bar is 0.5 µm. (E) Invasion efficiency (percentage of the inoculum that was internalized) of *P. alcalifaciens* WT, Δ*invA*_1a_, Δ*invA*_1a_ carrying pGEN-MCS (empty vector) or pGEN-*invA*_1a_ (complemented strain), and Δ*invA*_1b_ bacteria into HCT116 cells, THP-1 cells and S2 cells at 1 h p.i. was quantified by gentamicin protection assay. Invasion efficiency of WT bacteria was set to 100% in each experiment. Mean ± SD from 3-4 independent experiments. Asterisks indicate data significantly different from WT or Δ*invA*_1a_ pGEN bacteria (p<0.05, ANOVA with Dunnett’s post-hoc test).

As *P. alcalifaciens* is known to colonize diverse hosts, we tested whether its entry into different types of eukaryotic cells was T3SS-dependent by using the gentamicin protection assay. *P. alcalifaciens* 205/92 WT and Δ*invA* deletion mutants were grown under conditions that induce the T3SS_1a_ (Figure 2C) prior to their addition to mammalian and insect cells. Internalized bacteria were enumerated at 1 h post-infection (p.i.) and invasion efficiency was calculated as the proportion of the bacterial inoculum that was internalized. Compared to wild type (WT) bacteria, Δ*invA*_1a_ bacteria were highly defective for entry into HCT116 human colonic epithelial cells, and *D. melanogaster* S2 cells, which have characteristics of fly hemocytes (Figure 3E). In macrophages derived from the human monocytic cell line, THP-1, there was no difference in the internalization efficiency of the three bacterial strains, in accordance with the phagocytic properties of these cells. By contrast, the invasion efficiency of Δ*invA*_1b_ bacteria was comparable to WT bacteria in HCT116, THP-1 and S2 cells (Figure 3E). Overall, we conclude that *P. alcalifaciens* T3SS_1a_ is required for efficient bacterial internalization into human IECs and insect cells, suggesting a role for the T3SS_1a_-dependent translocation of type III effectors in this entry process.

### Intracellular replication of P. alcalifaciens

We next investigated whether *P. alcalifaciens* 205/92 can survive and replicate intracellularly in mammalian and insect cells. In HCT116 epithelial cells, there was a ∼3-fold increase in recoverable CFUs for WT bacteria over a 12 h time course (Figure 4A, left panel), with the greatest net increase in intracellular proliferation occurring between 1 and 4 h p.i. (Figure 4A, left panel). We confirmed the intracellular proliferation of WT bacteria within epithelial cells using an inside/outside assay in conjunction with fluorescence microscopy. HCT116 cells were infected with *P. alcalifaciens* carrying a plasmid constitutively expressing *dsRed* (pGEN-DsRed.T3) and anti-*P. alcalifaciens* antibodies were used to probe for extracellular bacteria in non-permeabilized cells. The number of intracellular WT bacteria increased from 1 h p.i. (mean of 1.7 bacteria/cell) to 8 h p.i. (mean of 2.92 bacteria/cell) (Figure 4A, middle panel). Replication of the Δ*invA*_1b_ mutant was indistinguishable from WT bacteria in HCT116 cells (Figure 4A, 4B). However, there was a progressive decrease in recoverable CFUs for Δ*invA*_1a_ bacteria by gentamicin protection assay (Figure 4A, left panel) and no evidence of bacterial proliferation at the single-cell level (mean of 1.6 bacteria/cell and 1.8 bacteria/cell at 1 h p.i. and 8 h p.i., respectively) (Figure 4A, middle panel). Negligible cytotoxicity was measured over the infection time course, irrespective of the infecting bacterial strain (Figure 4A, right panel).

**Figure 4.**
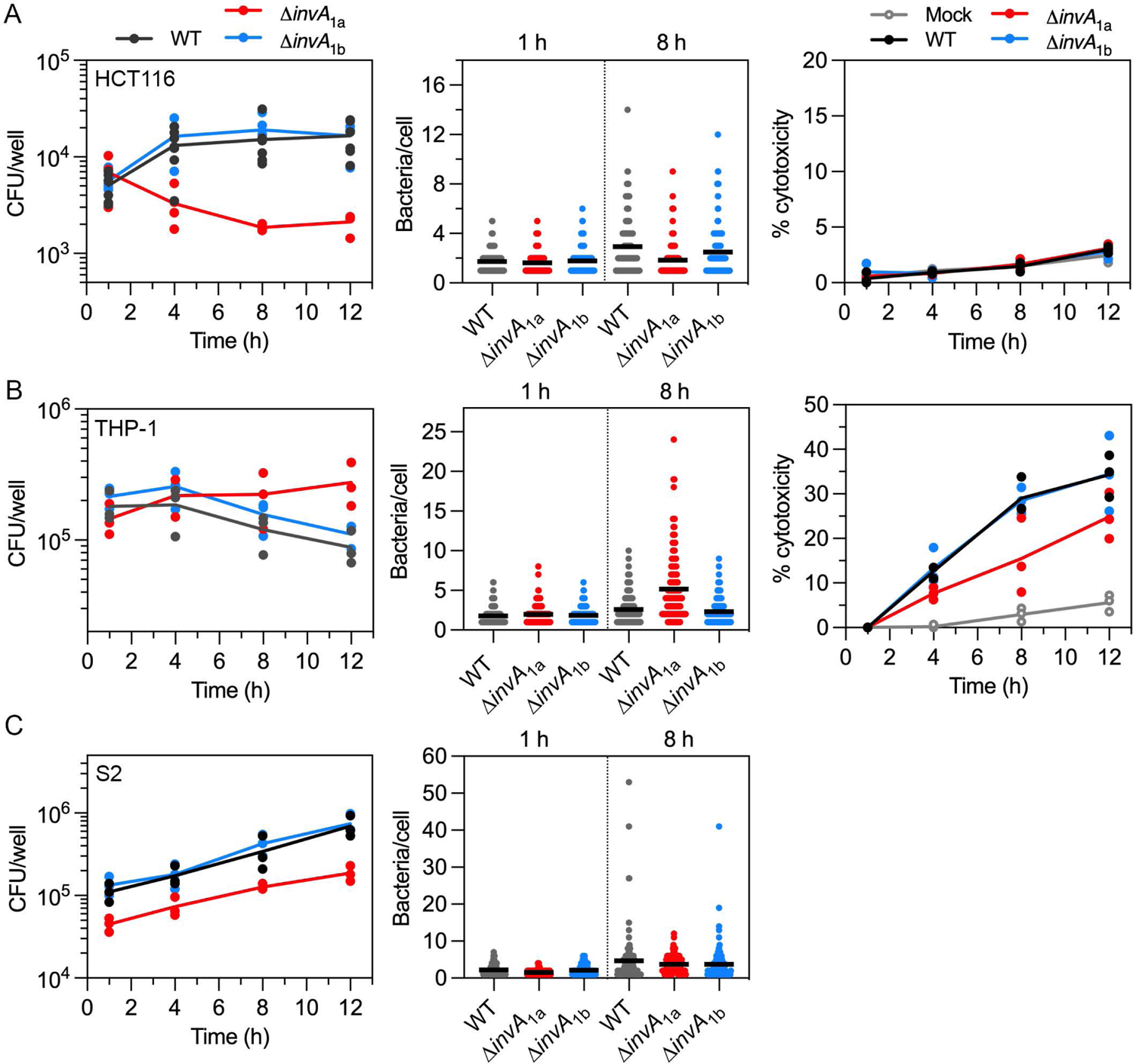
Intracellular replication of *P. alcalifaciens* in eukaryotic cells. (A) Infection of HCT116 epithelial cells. Intracellular proliferation of WT, Δ*invA*_1a_ and Δ*invA*_1b_ bacteria was quantified by gentamicin protection assay (left panel, n=3 independent experiments, each dot represents the mean from one experiment) and fluorescence microscopy (middle panel, each dot represents the number of bacteria in one cell, horizontal bar indicates the mean of 3 independent experiments). For microscopy experiments, WT, Δ*invA*_1a_ and Δ*invA*_1b_ bacteria were carrying pGEN-DsRed.T3. Inside/outside staining was used to distinguish intracellular from extracellular DsRed-labelled bacteria. Cell death was measured by LDH release into the supernatants (right panel, n=3 independent experiments, each dot represents the mean from one experiment). Percent cytotoxicity was calculated by normalizing to maximal cell death (1% (v/v) Triton X-100 lysis). (B) Infection of THP-1 macrophages. Intracellular proliferation of WT, Δ*invA*_1a_ and Δ*invA*_1b_ bacteria was quantified by gentamicin protection assay (left panel, n=3 independent experiments) and fluorescence microscopy (middle panel, n=3 independent experiments). Cytotoxicity was measured using LDH release assay (right panel, n=3 independent experiments). (C) Infection of S2 cells. Intracellular proliferation of WT, Δ*invA*_1a_ and Δ*invA*_1b_ bacteria was quantified by gentamicin protection assay (left panel, n=3 independent experiments) and fluorescence microscopy (middle panel, n=2 independent experiments).

The patterns of intracellular proliferation for WT bacteria and the deletion mutants were different in human macrophages as compared to human IECs. Using the gentamicin protection assay, there was a decrease in recoverable CFUs for WT bacteria over a 12 h time course of infection in THP-1 cells (Figure 4B, left panel). Incongruently, there was a slight increase in the mean number of WT bacteria/cell at the single-cell level, from a mean of 1.8 bacteria/cell at 1 h p.i. to 2.6 bacteria/cell at 8 h p.i., (Figure 4B, middle panel). However, high levels of cell death/cytotoxicity during WT infection, up to 35% of the monolayer by 12 h p.i. (Figure 4B, right panel), likely account for the overall decrease in viable CFUs as assessed by the gentamicin protection assay. Δ*invA*_1b_ bacteria showed a similar profile to WT bacteria for CFUs, number of bacteria/cell, and cytotoxicity induction kinetics (Figure 4B). By contrast, there was a net increase in Δ*invA*_1a_ bacteria over time as measured by CFUs and the number of bacteria per cell (mean of 2.0 and 5.2 bacteria/cell at 1 h p.i. and 8 h p.i., respectively), indicating replication of Δ*invA*_1a_ bacteria in THP-1 cells (Figure 4B). However, significantly less LDH was released into the culture supernatants, suggesting that the overall increase in viable Δ*invA*_1a_ bacteria was explained, in part, by decreased THP-1 cell death/detachment of infected cells compared to WT infections (Figure 4B).

Upon *P. alcalifaciens* infection of *D. melanogaster* S2 cells, there was an overall increase in intracellular CFUs and the mean number of bacteria per cell for WT, Δ*invA*_1a_ and Δ*invA*_1b_ bacteria (Figure 4C). Notably, the replication level of WT bacteria over 12 h (gentamicin protection assay) and 8 h (fluorescence microscopy) was higher in insect cells than in mammalian cells. The colorimetric assay for LDH release cannot be used with S2 cells (Walker et al., 2013) so we were unable to assess host cell cytotoxicity for *Providencia*-S2 cell infections. Collectively, these data indicate that *P. alcalifaciens* 205/92 can replicate intracellularly in mammalian and insect cells and the contribution of T3SS_1a_ to bacterial proliferation and induction of host cell death is both host- and cell-type dependent.

### Intracellular expression kinetics of T3SS_1a_ and T3SS_1b_

To follow *P. alcalifaciens* gene expression after internalization into mammalian and insect cells, we infected HCT116 and S2 cells with WT bacteria carrying plasmid-borne P*invF*_1a_-*luxCDABE* or P*prgH*_1a_-*luxCDABE* as T3SS_1a_ reporters, or P*invF*_1b_-*luxCDABE* or P*prgH*_1b_-*luxCDABE* as T3SS_1b_ reporters. At various times p.i., infected monolayers were collected, lysed and luminescence associated with intracellular bacteria quantified in a plate reader. We found that the intracellular expression kinetics of *P. alcalifaciens* genes encoded in T3SS_1a_ are comparable to those in *S*. Typhimurium T3SS1 (Ibarra et al., 2010). Specifically, we observed a rapid down-regulation of *prgH*_1a_ and *invF*_1a_ gene transcription, such that by 6 h p.i., bacterial luminescence was equivalent to the promoterless *luxCDABE* plasmid backbone, pFU35 (Figure 5). Transcription of *prgH*_1b_ and *invF*_1b_ was not observed over the time course indicating that T3SS_1b_-associated genes are not induced intracellularly in mammalian or insect tissue culture cells, at least up to 6 h p.i.

**Figure 5.**
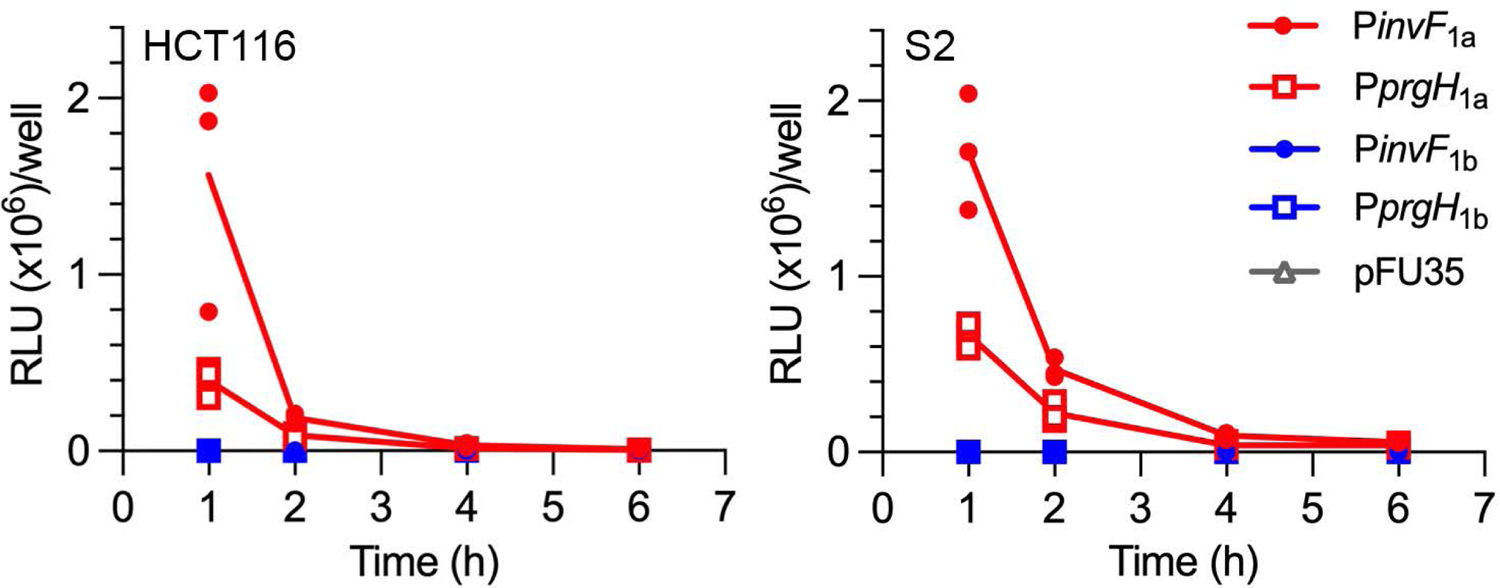
Rapid down-regulation of T3SS_1a_ after bacterial internalization. HCT116 (left panel) or S2 (right panel) cells were infected with *P. alcalifaciens* WT bacteria carrying one of the following transcriptional reporter plasmids: P*invF*_1a_-*luxCDABE*, P*prgH*_1a_-*luxCDABE*, P*invF*_1b_-*luxCDABE*, P*prgH*_1b_-*luxCDABE* or the empty vector control (pFU35). At the indicated times, infected monolayers were collected, lysed and the luminescence associated with internalized bacteria measured in a plate reader. Line indicates the mean of three independent experiments (each symbol represents data from one experiment).

### P. alcalifaciens colonizes the cytosol of eukaryotic cells

Our earlier work on the SipC/IpaC family of type III translocators showed that these proteins have differential destabilizing activities on bacteria-containing vacuole membranes (64). For example, replacing SipC in *S*. Typhimurium with its ortholog from either of the professional cytosolic pathogens, *S. flexneri* (IpaC) or *Chromobacterium violaceum* (CipC), enables *S*. Typhimurium to lyse its internalization vacuole more efficiently. We used this gene swapping strategy to predict the intracellular niche of *P. alcalifaciens* 205/92. We have previously shown that *P. alcalifaciens* SipC_1a_ can partially complement for the internalization defect of a *S*. Typhimurium Δ*sipC* mutant (Δ*sipC*::*sipC*_1a_) into non-phagocytic cells (50). Using the chloroquine resistance assay (65, 66), which determines the proportion of internalized bacteria that are present in the cytosol, we found a significantly increased proportion of *S*. Typhimurium Δ*sipC*::*sipC*_1a_ bacteria in the cytosol of HCT116 epithelial cells and J774A.1 macrophages compared to *S*. Typhimurium WT (Figure 6A). Notably, the level of nascent vacuole lysis for Δ*sipC*::*sipC*_1a_ and Δ*sipC*::*cipC* bacteria was comparable (Figure 6A). From these results, we speculated that *P. alcalifaciens* is a cytosolic bacterium, like *C. violaceum* (64).

**Figure 6.**
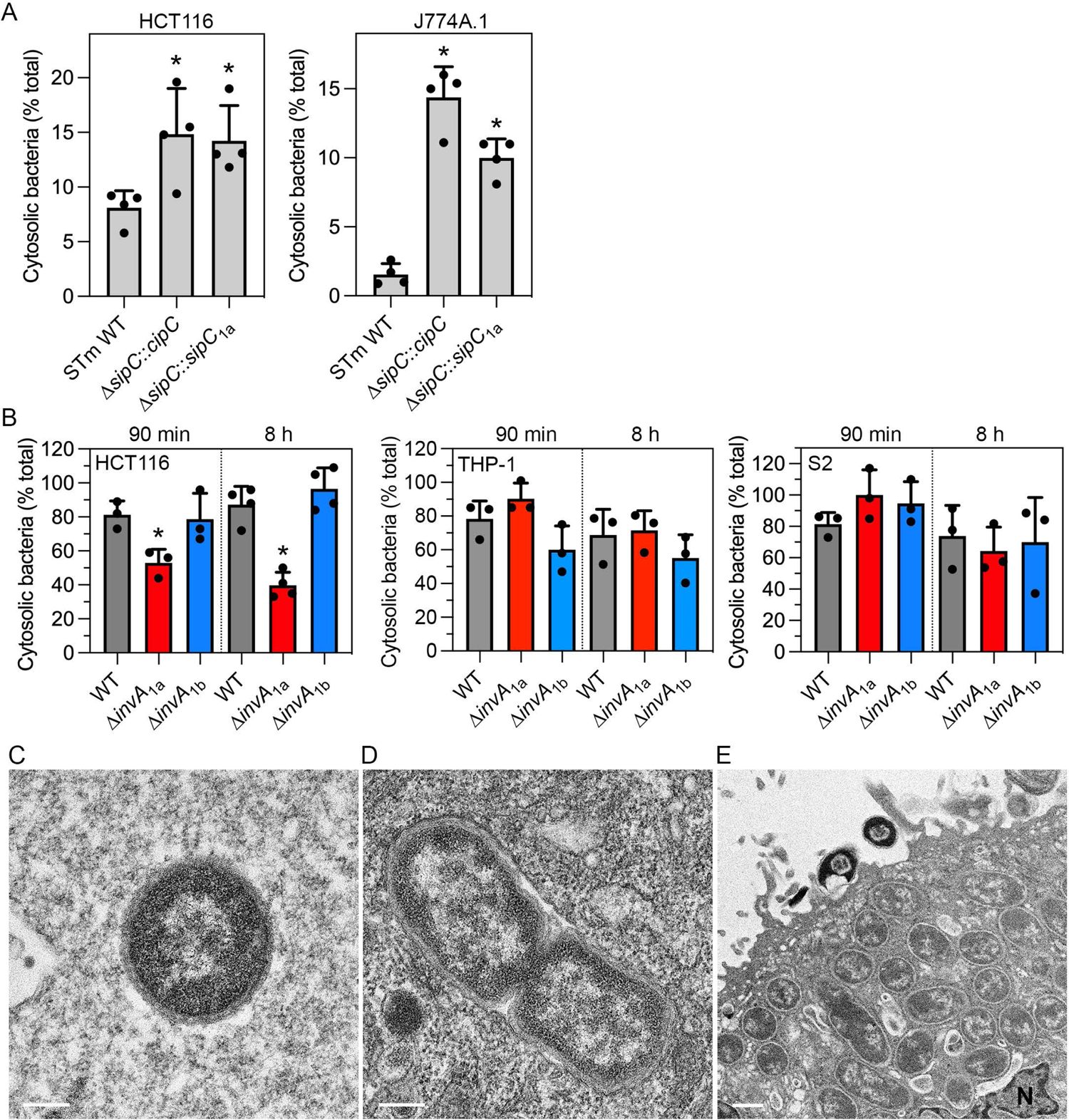
*P. alcalifaciens* lyses its internalization vacuole, then replicates in the cytosol. (A) HCT116 (left panel) or J774A.1 (right panel) cells were infected with *S*. Typhimurium (STm) WT, Δ*sipC*::*cipC* (chromosomal replacement of *sipC* with *C. violaceum cipC*) or Δ*sipC*::*sipC*_1a_ (chromosomal replacement of *sipC* with *P. alcalifaciens sipC*_1a_) bacteria and the proportion of internalized bacteria present in the cytosol at 90 min p.i. was assessed by CHQ resistance assay. Mean ± SD from 4 independent experiments. Asterisks indicate data significantly different from STm WT bacteria (p<0.05, ANOVA with Dunnett’s post-hoc test). (B) HCT116 (left panel), THP-1 (middle panel) and S2 (right panel) cells were infected with *P. alcalifaciens* 205/92 WT, Δ*invA*_1a_ and Δ*invA*_1b_ bacteria and the proportion of cytosolic bacteria at 90 min and 8 h p.i. was determined by CHQ resistance assay. Mean ± SD from 3-4 independent experiments. Asterisks indicate data significantly different from WT bacteria (p<0.05, ANOVA with Dunnett’s post-hoc test). (C and D) Representative TEM images of cytosolic (C) and vacuolar (D) *P. alcalifaciens* in epithelial cells at 90 min p.i. Scale bars are 200 nm. (E) Representative TEM image of *P. alcalifaciens* replicating in the cytosol of an epithelial cell at 4 h p.i. Scale bar is 500 nm, N=nucleus.

We applied the CHQ resistance assay to *Providencia*-infected HCT116, THP-1 and S2 cells to unequivocally determine their intracellular replication niche. At 90 min p.i., ∼80% of WT bacteria were present in the cytosol in all three cell types (Figure 6B). Therefore, lysis of the nascent bacteria-containing vacuole is fast and efficient. This level of cytosolic presence was sustained at later times (>70% cytosolic bacteria at 8 h p.i.), indicating bacterial replication (Figure 4) occurs in the cytosol of eukaryotic cells (Figure 6B). There was no significant difference in the proportion of cytosolic WT and Δ*invA*_1b_ bacteria in any cell type (Figure 6B). However, a lower proportion of Δ*invA*_1a_ bacteria were present in the cytosol of HCT116 cells at 90 min and 8 h p.i. (Figure 6B), indicating a defect in nascent vacuole lysis in IECs. This dependence on T3SS_1a_ for vacuole lysis was not observed in human macrophages or insect cells, however (Figure 6B).

We used TEM analysis to independently assess the intracellular locale of *P. alcalifaciens* WT in eukaryotic cells. Most bacteria were free in the cytosol of epithelial cells by 90 min p.i. (59% cytosolic in HeLa cells, n=49; 64% cytosolic in HCT116 cells, n=11) (Figure 6C). In some cases, an intact vacuolar membrane was observed around bacteria in epithelial cells at 90 min p.i. (Figure 6D). By 4 h p.i., bacterial replication in the cytosol of epithelial cells was evident (Figure 6E). We also observed bacteria that remain extracellular, seemingly firmly attached to the surface of epithelial cells, that were killed by gentamicin (Figure 6E). Like epithelial cells, most intracellular WT bacteria were found in the cytosol, without any obvious surrounding membrane, in THP-1 macrophages (83% cytosolic, n=18) and S2 cells (93% cytosolic, n=43) by 90 min p.i. Collectively, these data support that *P. alcalifaciens* 205/92 rapidly escapes from its internalization vacuole, then replicates in the cytosol of mammalian and insect cells.

### Inflammasome activation by P. alcalifaciens

Inflammasomes are cytosolic innate immune sensors that play a key role in restricting bacterial infections and epithelial-intrinsic inflammasomes mediate protective responses against intestinal pathogens such as *S*. Typhimurium (67–69) and *C. rodentium* (70). Considering *P. alcalifaciens* can invade mammalian cells (Figures 3 and 4) and replicate in the cytosol (Figure 6), we speculated it would activate human IEC canonical (caspase-1) and/or non-canonical (caspase-4) inflammasomes. We have previously shown that the activation status of murine IECs affects the contribution of non-canonical and canonical inflammasomes to host defense (71). We therefore considered that inflammasome activation in human IECs might also be similarly affected. In agreement with previous reports in HT-29 colonic epithelial cells (72–74), *CASP1* mRNA was significantly up-regulated by IFNγ treatment in HCT116 cells (Figure 7A). A robust increase in pro-caspase-1 and moderate increase in pro-caspase 4 levels were detected upon IFNγ treatment (Figure 7B). Therefore, with IFNγ priming in conjunction with *CASP1*^-/-^ and *CASP4*^-/-^ knockout (KO) cells, we can investigate whether *P. alcalifaciens* 205/92 activates canonical and non-canonical inflammasomes in IECs.

**Figure 7.**
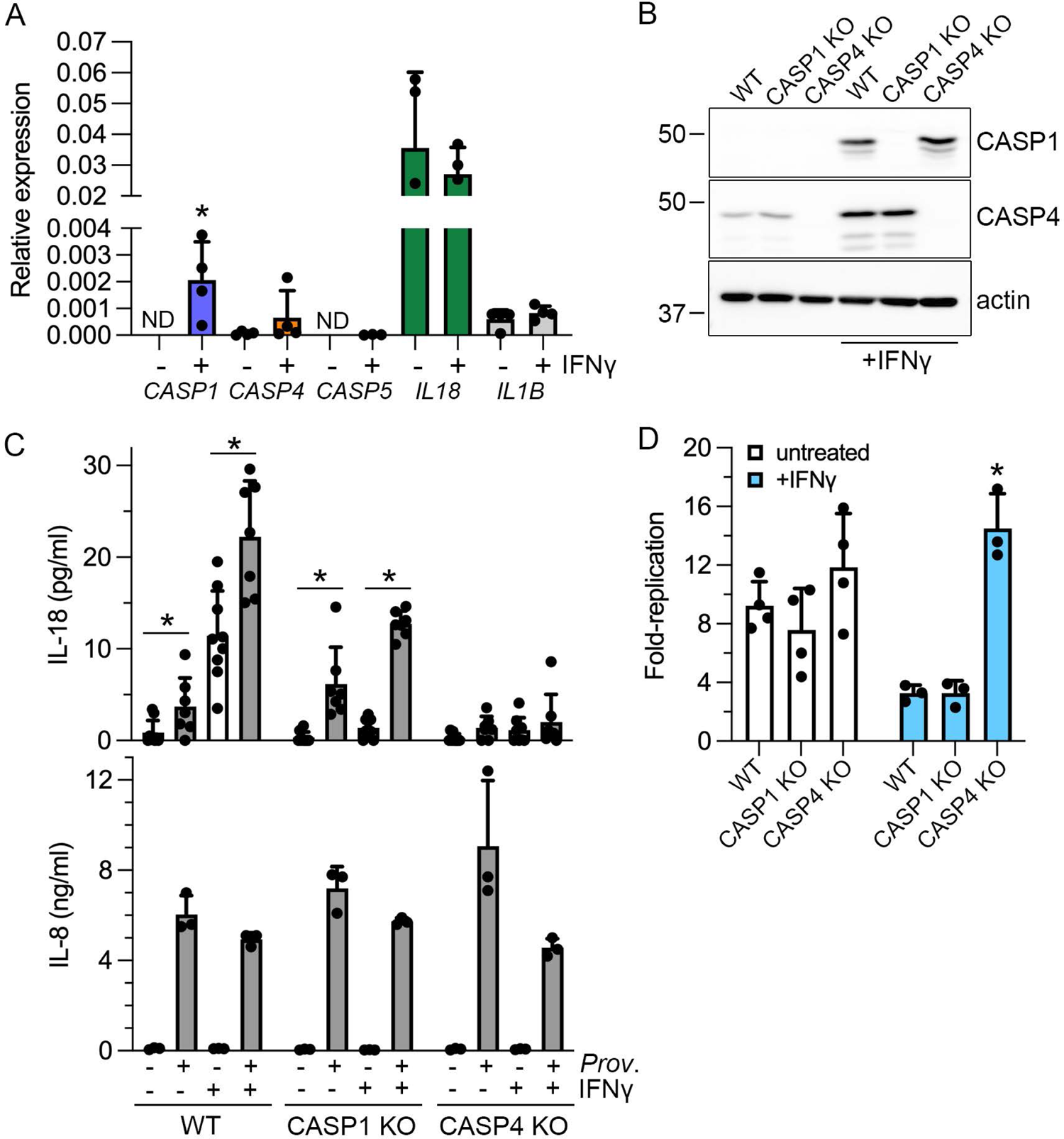
*P. alcalifaciens* 205/92 activates the non-canonical inflammasome in IECs. (A) HCT116 cells were left untreated or treated with 50 ng/ml IFNγ for 16-18 h. mRNA expression of *CASP1*, *CASP4*, *CASP5*, *IL18* and *IL1B* relative to the reference gene *RPLP0* was measured by qPCR (expressed as 2^-ΔCq^). n=3-4 independent experiments. Asterisk indicates data significantly different from untreated cells, p<0.05, Student’s t-test. (B) Immunoblot analysis of caspase-1, caspase-4 and β-actin (loading control) in HCT116 WT, *CASP1*^-/-^ (CASP1 KO) or *CASP4*^-/-^ (CASP4 KO) cells, left untreated or treated with IFNγ for 16-18 h. (C) IL-18 (upper panel) and IL-8 (lower panel) release into supernatants from mock-infected or *P. alcalifaciens*-infected HCT116 WT, CASP1 KO or CASP4 KO cells at 16 h p.i. was quantified by ELISA. n≥8 independent experiments. Cells were treated with IFNγ (50 ng/ml) for 16-18 h prior to infection where indicated (+). Asterisks indicate significantly different data, p<0.05, Student’s t-test. (D) Bacterial replication in HCT116 WT, CASP1 KO and CASP4 KO cells was assessed by gentamicin protection assay. Fold-replication is CFU_10h_/CFU_1h_. Asterisk indicates data significantly different from HCT116 WT, p<0.05, ANOVA with Dunnett’s post-hoc test. n=3-4 independent experiments.

Processing and secretion of two cytokines, interleukin (IL)-18 and IL-1β, is dependent on inflammasome activation. Despite encoding *IL1B* mRNA ((75); Figure 7A), IL-1β is not secreted by human IECs upon bacterial infection ((76); our unpublished results). We therefore used IL-18 secretion as a readout of inflammasome activation. IL-8 release was a control for an inflammasome-independent cytokine. Upon *Providencia* infection of WT HCT116 cells, IL-18 was detected in the cell culture supernatant at 16 h p.i. in both untreated and IFNγ-primed cells (Figure 7C). The infection-dependent potentiation of IL-18 release from both untreated and IFNγ-primed cells was statistically significant (Figure 7C). The magnitude of increase in IL-18 release after *P. alcalifaciens* infection of *CASP1*^-/-^ KO cells was comparable to that of WT HCT116 cells, with and without IFNγ treatment (Figure 7C). By contrast, IL-18 release was abrogated from infected *CASP4*^-/-^ KO cells, irrespective of IFNγ-priming (Figure 7C). Infection-induced IL-8 secretion was comparable regardless of priming status or cell genotype (Figure 7C). These data indicate that *P. alcalifaciens* 205/92 stimulates IL-18 secretion from human IECs in a caspase-4-dependent manner i.e. via activation of the non-canonical inflammasome. Previous studies in human IECs have shown that caspase-4 is critical for limiting *S*. Typhimurium replication (67, 76, 77). To test whether IEC intrinsic inflammasomes (caspase-1 and/or caspase-4) influence intracellular replication of *P. alcalifaciens* 205/92, we compared bacterial replication over a 10 h timeframe. In unprimed IECs, there was no significant difference in *P. alcalifaciens* replication in WT, *CASP1*^-/-^ or *CASP4*^-/-^ HCT116 cells (9.2-, 7.6- and 11.9-fold, respectively; Figure 7D). IFNγ-priming efficiently restricted bacterial replication in WT and *CASP1*^-/-^ HCT116 cells (3.3- and 3.3-fold, respectively) but not *CASP4*^-/-^ cells. Rather, robust bacterial replication was still observed in *CASP4*^-/-^ IECs upon IFNγ-priming (14.5-fold; Figure 7D). Therefore, caspase-4, but not caspase-1, restrict the intracellular proliferation of *P. alcalifaciens* in human IECs.

### Infection models

We used the bovine ligated intestinal loop model, which we have previously used to study another enteric pathogen, *S. enterica* serovar Typhimurium (78), to assess the enteropathogenicity of *P. alcalifaciens* 205/92. Bacteria (∼10^9^ CFU) were injected into each loop and loops were collected at 12 h post-inoculation. Most bacteria remained extracellular, with only 10^7^ WT bacteria being tissue-associated (Figure 8A). Compared to the LB control, infection with *P. alcalifaciens* 205/92 did not promote fluid accumulation in the calf model (Figure 8B), a proxy for intestinal secretory responses. Infection-induced inflammatory changes were assessed by histological evaluation of hematoxylin- and eosin-stained sections of tissue samples. Of the six criteria scored – polymorphonuclear infiltration to the lamina propria, submucosal edema, epithelial damage, villus blunting, crypt abscess and cell death – only epithelial damage was increased with statistical significance in infected tissues (p=0.04, Student’s t-test; Figure 8C, 8D, 8E). In this infection model, we did not observe any difference in fluid or tissue colonization (Figure 8F), fluid accumulation or local inflammatory responses (results not shown) when comparing infection with WT, Δ*invA*_1a_ or Δ*invA*_1b_ bacteria at 2 h or 8 h p.i.

**Figure 8.**
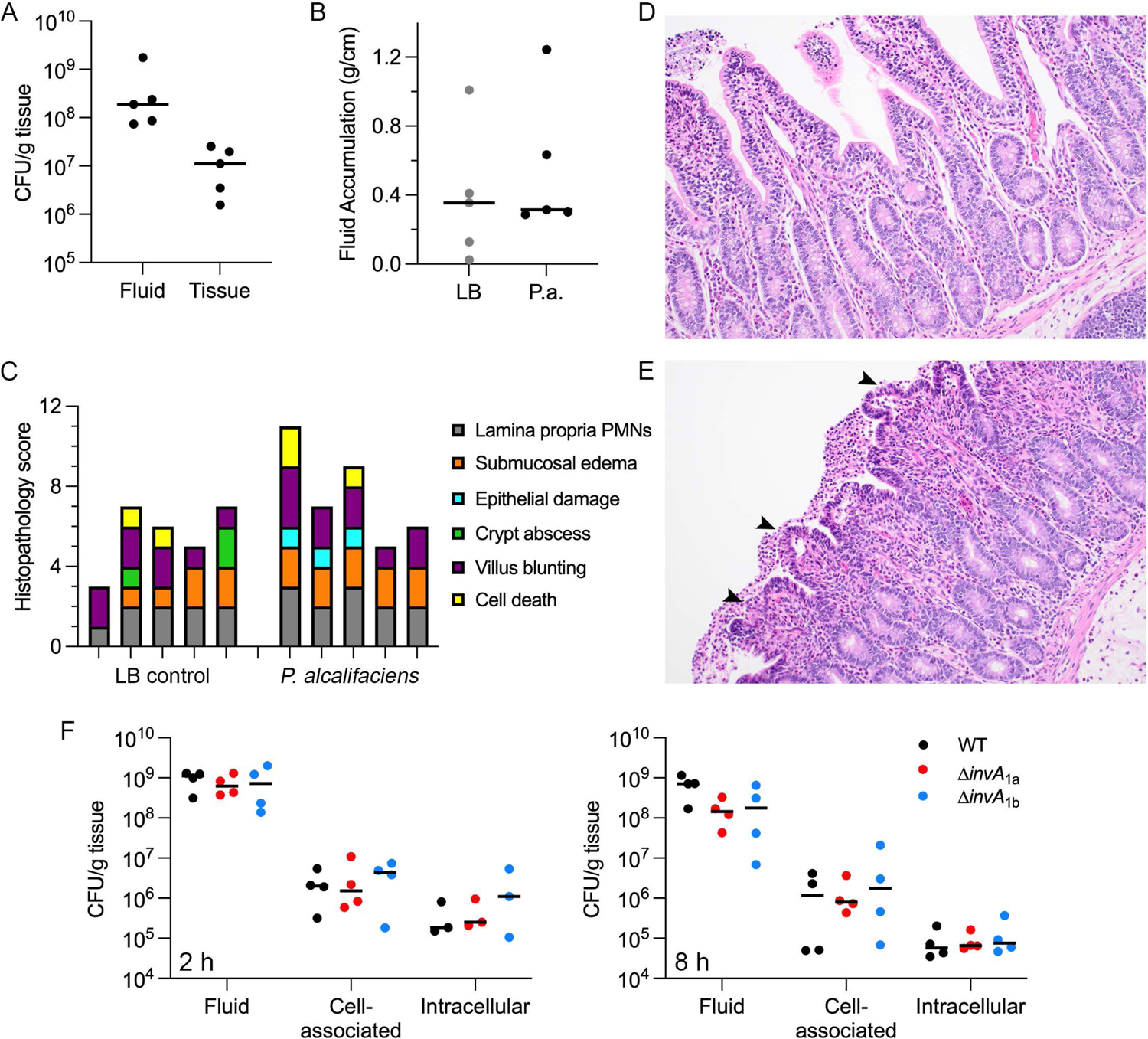
*P. alcalifaciens* 205/92 infection of bovine ligated intestinal loops. Bovine ligated intestinal loops were inoculated with LB broth (control) or *P. alcalifaciens* 205/92 (∼10^9^ CFU) resuspended in LB broth. (A) Recovery of WT bacteria from the fluid and tissue after 12 h. Each dot represents data from one calf. Means are indicated. (B) Secretory responses in intestinal loops after infection with WT bacteria (P.a.) or LB control (LB) for 12 h. Each dot represents data from one calf. Means are indicated. (C) Pathological scores from hematoxylin- and eosin-stained loop tissues sampled 12 h after inoculation with LB broth or WT bacteria. *P. alcalifaciens* infection led to increased epithelial damage in the intestine (p=0.040, Student’s t-test). Each bar represents one calf. (D) and (E) Representative images (20x objective) of hematoxylin- and eosin-stained sections of intestinal tissue from loops inoculated with LB broth alone (D) or WT bacteria (E). Sites of epithelial damage are indicated by arrowheads. (F) Recovery of WT, Δ*invA*_1a_ and Δ*invA*_1b_ bacteria from the fluid and tissue (cell-associated or intracellular) after 2 h (left panel) and 8 h (right panel). Each dot represents the average of 2-3 loops per calf (2 h) or one loop per calf (8 h). Means are indicated.

As an alternative infection model, we used *D. melanogaster* to test whether T3SS1a or T3SS1b from *P. alcalifaciens* 205/92 were required for virulence. In insect hosts, *P. alcalifaciens* is highly virulent. In earlier studies, infection with 10^3^-10^4^ *P. alcalifaciens* Dmel, a strain originally isolated from wild *D. melanogaster*, causes mortality in 99% of flies within 2 days (13). In a separate study, all flies were dead within 27 h after an infection dose of ∼3750 CFU *P. alcalifaciens* DSM30120 (79). Of note, the genomes of Dmel and DSM30120 encode for T3SS_1b_ but not T3SS_1a_ (13)(Figure 1C, Table S1). In our initial tests with *P. alcalifaciens* 205/92, an infection dose of ∼300 CFU proved too high, with rapid killing of all flies between 30-40 h (results not shown). When we reduced the infection dose to 30 CFU, all flies succumbed to infection with WT bacteria within 45-68 h (Figure 9). Virulence of *P. alcalifaciens* Δ*invA*_1a_ bacteria was comparable to WT bacteria at an infection dose of 30 CFU (Figure 9, p=0.83) whereas Δ*invA*_1b_-infected flies showed an extended time to death (Figure 9, p<0.0001). *P. alcalifaciens*-induced mortality was dose-dependent; at a lower infectious dose of WT bacteria (10 CFU), 5-6% of flies survived for up to 93 h, when we stopped monitoring survival (Figure 9). Compared to WT bacteria, we observed a prolonged time to death, and decreased mortality, during infection with Δ*invA*_1a_ and Δ*invA*_1b_ bacteria at a dose of 10 CFU (Figure 9). Both survival curves were significantly different from infection with WT bacteria (p=0.0019 for Δ*invA*_1a_ and p<0.0001 for Δ*invA*_1b_), indicating that flies are less susceptible to infection with these gene deletion mutants. Overall, our results demonstrate two points: (1) that flies are highly susceptible to infection with *P. alcalifaciens* 205/92 and, (ii) that T3SS_1a_ and T3SS_1b_ are both virulence determinants in an insect host.

**Figure 9.**
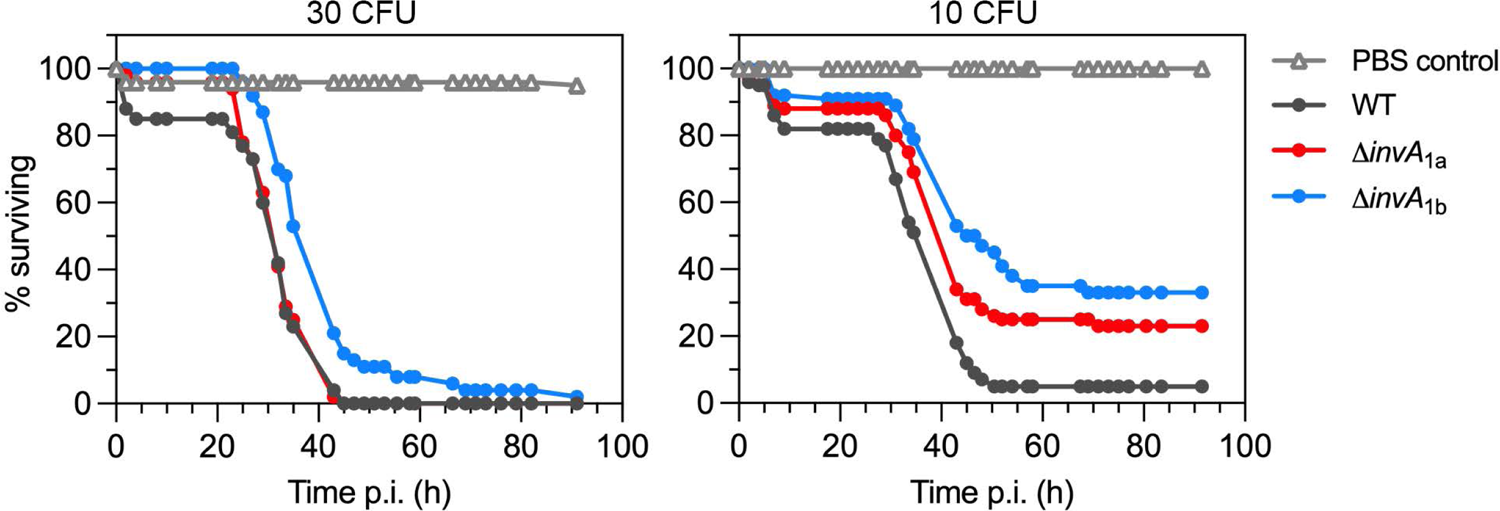
T3SS_1b_, and to a lesser extent T3SS_1a_, are virulence determinants in insects. *D. melanogaster* were inoculated with *P. alcalifaciens* 205/92 WT, Δ*invA*_1a_ or Δ*invA*_1b_ bacteria, or PBS, and survival was monitored. Results are from an infectious dose of ∼30 CFU/fly (left panel) or ∼10 CFU/fly (right panel) and are representative of 2-3 independent experiments. >50 flies were inoculated with PBS or each strain in each experiment.

## Discussion

T3SS play a central role in cell-cell interactions between bacteria and eukaryotes, irrespective of whether bacteria are pathogens, mutualists or commensals (80). The high prevalence of T3SS in *Providencia* spp. suggests this molecular system is important for their colonization of specific ecological niches. Up to nine T3SS families have been identified (81) and both T3SSs of *P. alcalifaciens* 205/92 belong to the Inv/Mxi-Spa family. Our earlier phylogenetic analysis of the translocator proteins in this family indicated that there are two sub-groups of T3SS (50). The first sub-group contains bacteria that colonize and cause disease in mammals i.e. *S. enterica*, *S. flexneri*, enteroinvasive *E. coli* (EIEC), and *Chromobacterium* spp. *P. alcalifaciens* T3SS_1a_ is also in this clade. The second sub-group, in which T3SS_1b_ belongs, contains bacteria that colonize diverse environments such as pathogens of fungi (*Pseudomonas gingeri*), plants (*Xanthomomas albilineans*), and insects (*Providencia sneebia*), endosymbionts of insects (*Sodalis glossinidius*) and opportunistic pathogens of humans (*Proteus mirabilis*). Notably, T3SS required for bacterial colonization of insect hosts are restricted to the second sub-group. For example, the SSR-2 T3SS from *S. glossinidus* enhances bacterial proliferation in insect cells (51), the Ysa T3SS aids in *Y. enterocolitica* replication in insect, and not mammalian, cells (82) and PSI-2 from *Pantoea stewartii* is required for persistence in the gut of its flea beetle vector (83). We have further shown that T3SS_1b_ is a *P. alcalifaciens* virulence factor in *D. melanogaster*. We predict that the T3SS_1b_ from other *Providencia* spp. i.e. *P. sneebia*, *P. rettgeri* and *P. vermicola* (19, 84, 85), will also promote insect infection. The differential contribution of T3SS_1a_ and T3SS_1b_ to vertebrate and invertebrate host colonization, respectively, likely explains their distinct responses to environmental cues. *S*. Typhimurium T3SS1 and *P. alcalifaciens* T3SS_1a_ genes are induced under similar *in vitro* conditions – late-log phase of growth, aeration, high salt (59, 86) – and rapidly down-regulated following bacterial internalization into eukaryotic cells (59) suggesting that they share commonalities in their regulatory networks. Indeed, *P. alcalifaciens* T3SS_1a_ encodes for orthologs of *hilA*, *invF* and *sicA* (Figure 2A), three of the main transcriptional regulators of T3SS1 activity in *S*. Typhimurium (87). *In vitro* growth conditions that induce T3SS_1b_ genes remain an enigma as altering growth phase, temperature, salt and pH were unsuccessful. We also did not observe T3SS_1b_ gene induction upon bacterial infection of mammalian (HCT116) or insect (S2) cells. However, the Δ*invA*_1b_ deletion mutant is attenuated in flies which indicates there are gene induction cues in the insect model that we have not been able to recapitulate in broth or tissue culture cells. Overall, our data supports the idea that *P. alcalifaciens* extended its host range from the natural environment to living organisms by acquiring these two divergent T3SS, each adapted to a target host class: *Mammalia* (T3SS_1a_) or *Insecta* (T3SS_1b_).

Gram-negative bacteria in the genus *Providencia* are considered opportunistic pathogens. *P. alcalifaciens* is part of the human gut microbiome (8) and has been isolated from the feces of healthy people and animals (21, 29, 39, 88), and wastewater treatment plants (89). *P. alcalifaciens* is also an enteropathogen (31, 32, 39, 40). Based on the available genome sequences and bacterial strain source details, we describe a strong link between the subset of *P. alcalifaciens* strains that harbor T3SS_1a_ (Figure 1) and cause diarrheal disease in dogs (e.g. strains 2019-04-29292-1-3 and 2019004029290-1-7; Table S1) and humans (e.g. strains 205/92, 2939/90 and RMID 1656011; Table S1). Therefore it seems that the horizontal transfer of genes encoding a second T3SS on the 128kb plasmid has contributed to the emergence of enteropathogenic *P. alcalifaciens* strains (44, 90). Also encoded on the 128kb plasmid of *P. alcalifaciens* 205/92 are homologs of type III effectors known to be bacterial virulence determinants in mammalian hosts, including SptP, EspG, EspM and SipA_1a_. SptP from *S. enterica* has two distinct functions. Its tyrosine phosphatase activity interferes with extracellular-regulated kinase (ERK) MAP kinase pathways in the host cell (91). It also acts a GTPase activating protein (GAP) for Rac1 and Cdc42 to reverse actin cytoskeletal changes that accompany bacterial entry into epithelial cells (92). The EspG protein family consists of EspG/EspG2 and VirA (93). EspG from enteropathogenic *E. coli* (EPEC) disrupts the host cell secretory pathway (94). VirA is required for efficient entry of *Shigella* into epithelial cells, and intra- and inter-cellular bacterial spread (95). The EspM/Map/IpgB2 family of effectors are found in EPEC, EHEC and *Shigella* and modulate actin dynamics (96). *S. enterica* SipA and *S. flexneri* IpaA are functional orthologs that induce a loss of actin stress fibers to promote bacterial entry into non-phagocytic cells (97, 98). In addition to its actin-binding role, SipA is necessary and sufficient for polymorphonuclear leukocyte (PMNs) migration across the intestinal epithelium in both *in vitro* and *in vivo* models of *S*. Typhimurium infection (99). Of interest, the molecular mass of SipA_1a_ from a subset of *P. alcalifaciens* strains (e.g. strains 205/92, 2939-90, 2019-04-28369-1-2, 2019-04-29292-1-3, 2019-04-29034-1-3, 2019-04-2920-1-7, 2019-04-27799-1-2 and 2019-04-3283-1-1; Figures 1 and 2) is unusually large compared to SipA from *S. enterica*, IpaA from *S. flexneri*, or CipA from *Chromobacterium violaceum*. This is due to an extended C-terminus (Figure 1B); the biological function of this SipA_1a_ domain is unknown. The chromosome of *P. alcalifaciens* 205/92 also encodes for homologs of known type III effectors including SteB and SteC from *S*. Typhimurium and a second SipA (SipA_1b_). SteB contributes to *S*. Enteritidis biofilm formation on plastics (100) but the role of SteB in *Salmonella*-eukaryotic cell interactions has not yet been deciphered (101). SteC is a kinase that promotes actin cytoskeleton reorganization around the *Salmonella*-containing vacuole p(102, 103). Theoretically, this type III effector repertoire would allow *P. alcalifaciens* to enter and proliferate within eukaryotic cells. We are currently investigating whether these are *bona fide* type III effectors and if there are additional type III effectors of *P. alcalifaciens*.

We report that *P. alcalifaciens* 205/92 rapidly and efficiently lyses its internalization vacuole in mammalian and insect cells, then proliferates in the eukaryotic cell cytosol. A previous study showed that *P. alcalifaciens* 101i/59, an invasive isolate, was enclosed within vacuoles and present in the cytosol of Caco-2 cells at 4-6 h p.i. using crystal violet staining (48). Another invasive isolate, 82A-5778, occupied vacuoles “in close proximity to the nuclear membrane” and the cytosol in Hep-2 cells (45). No quantification of vacuolar versus cytosolic residence was provided in these two studies, however. We speculate that all *P. alcalifaciens* isolates that harbor T3SS_1a_ adopt a cytosolic lifestyle inside eukaryotic cells. We further identified mammalian cell responses that are indicative of host sensing of cytosolic bacteria, namely activation of the caspase-4 (non-canonical) inflammasome, followed by inflammatory cytokine release upon *P. alcalifaciens* infection of IECs. Caspase-4 also limits the cytosolic proliferation of *P. alcalifaciens*. There is no evidence for caspase-1 dependent host responses to *P. alcalifaciens* in colonic epithelial cells, in line with studies of other enteric pathogens in human IECs (76, 77).

In addition to inflammasome activation, ubiquitin-mediated autophagy is also an important host innate defense against cytosolic bacteria. Here we describe that nascent vacuole lysis is partially dependent on T3SS_1a_ in IECs but not human macrophages or insect cells (Figure 6), indicating host and cell type differences in *P. alcalifaciens* disruption of the surrounding vacuole membrane. Failure of Δ*invA*_1a_ bacteria to efficiently lyse the nascent vacuole in IECs results in bacterial killing (Figure 4). This hints that type III effectors are involved in disrupting the internalization vacuole membrane and preventing the autophagic recognition of *P. alcalifaciens* in human IECs. Orthologs of type III effectors known to interfere with autophagic recognition (VirA and IcsB in *S. flexneri*, SopF in *S*. Typhimurium, TssM in *Burkholderia* spp.) are not present in the genome of *P. alcalifaciens* 205/92, however. Other known mechanisms used by bacteria to deflect targeting by selective autophagy are LPS and O-antigen modifications (104, 105), enzymatic modification of bacterial outer membrane proteins (105) and bacterial phospholipase manipulation of host cell phospholipids (106–108). Our research efforts are currently directed towards defining how *P. alcalifaciens* mediates vacuolar escape and resists autophagic detection in mammalian cells.

Prior to our work, rabbits were the sole animal model used to study the enteropathogenesis of *P. alcalifaciens* infection. Murata *et al*. (2001) studied the secretory and inflammatory responses to *P. alcalifaciens* clinical isolates from a foodborne outbreak in Japan in rabbit ileal loops (31). Histopathological analysis of loops infected with one of these isolates (RIMD1656011) showed extensive mucosal inflammation including neutrophil infiltration within the lamina propria and distortion of the villus architecture. Seven isolates caused a moderate level of fluid accumulation after 20 h. By contrast, three isolates from diarrheal patients, one from Bangladesh (2939/90) and two from Australia (F90-2004, R90-1475), failed to induce fluid accumulation in the rabbit ileal loop assay after 20 h (40). Likewise, four *P. alcalifaciens* isolates from a foodborne outbreak in Kenya did not induce fluid accumulation in rabbit ileal loops over 18 h in a third study (33). Here, in a calf model of infection, fluid accumulation was not observed in intestinal loops exposed to *P. alcalifaciens* 205/92 for 12 h. Histopathological analysis of infected loops also did not show indicators of an acute inflammatory reaction, but mild epithelial damage was observed. Collectively, these studies suggest that either *P. alcalifaciens* causes non-inflammatory diarrhea, or intestinal loops in rabbits and calves are not an appropriate animal model to study the enteropathogenic properties of this bacterium. It should be noted, however, that the removable intestinal tie adult rabbit diarrhea (RITARD) model, which was developed in the early 1980’s to study *Vibrio cholerae* and enterotoxigenic *E. coli* enteric infections (109), supports the diarrheagenic nature of *P. alcalifaciens* clinical isolates (40).

*Providencia* spp. have been isolated from the hemolymph of wild caught *D. melanogaster* (13) and as part of the gut microbiome in *Bactrocera dorsalis*, the Oriental fruit fly (110), but it is not known if these bacteria are intracellular in insects. Fruit flies are exquisitely sensitive to infection by *P. alcalifaciens* (our results; (13, 79)). In *D. melanogaster* infections, *P. alcalifaciens* proliferates rapidly, reaching bacterial loads of ∼10^7^ - 10^8^ CFU per fly by 20-32 h p.i. (13, 79). Flies die shortly after (our results; (13, 79)). On the host side, Imd-dependent antimicrobial peptides and hemocyte-derived reactive oxygen species are the major branches of immunity that are important for fighting infection with *P. alcalifaciens* (79). In a forward genetics screen using a *P. alcalifaciens* DSM30120 transposon mutant library, lipopolysaccharide (LPS) and lipoprotein mutants showed reduced virulence in *D. melanogaster* (79). No bacterial mutants in toxins or T3SSs were hit in this screen (79). By contrast, we identified that T3SS_1b_, and to a lesser extent T3SS_1a_, are virulence factors in *D. melanogaster* (Figure 9), which is the first report of *P. alcalifaciens* T3SSs being virulence determinants in insects. We believe that the different results might be explained by the much lower infection dose used in our study (10-30 CFU versus 1500 CFU). Considering their high prevalence among *Providencia* spp. genomes (49), our results suggest that T3SSs are a widespread pathogenicity factor for invertebrate colonization by members of this genus. Furthermore, insects have a well-known role in transmitting clinically relevant pathogens, and *Providencia* spp. are often resistant to multiple antimicrobials, so studying the insect carriage of bacteria such as *P. alcalifaciens* may provide information about the role that flies play in the maintenance, and transmission, of antimicrobial resistance.

While historically viewed as opportunistic pathogens, the *Morganella*-*Proteus*-*Providencia* (MPP) group of organisms (*Morganella morganii*, *Proteus vulgaris,* and *Providencia* species) are increasingly being recognized as emerging causes of multidrug-resistant infections because of inducible chromosomal β-lactamases and a propensity to acquire other resistance determinants. The genomic plasticity of *Providencia* spp. is noteworthy, as it can be seen by the varied lifestyles of different species and strains, ranging from commensal residents of the gastrointestinal tract to assorted pathogens that promote intestinal or extraintestinal illnesses with different clinical consequences. Whole-genome sequencing has provided considerable information about the genomes of non-pathogenic and pathogenic *P. alcalifaciens* and what genes might contribute to the pathogenicity of this bacterial species (49, 90). From our work, and that of others, it is clear that some *P. alcalifaciens* strains have gained the ability to enter and proliferate within mammalian cells, cause damage to the gut epithelium and subsequent diarrheal disease. Acquisition of an “invasion-associated” plasmid drove this evolutionary leap. Altogether, our work argues that the importance of *P. alcalifaciens* as a *bona fide* enteropathogen should not be ignored and supports its inclusion into systematic surveillance programs.

## Materials and Methods

### Bacterial strains and plasmid construction

*P. alcalifaciens* 205/92 (Tet^R^) served as the wild-type (WT) strain and background for deletion mutants in this study. It was originally isolated from the stool of a 12-year-old boy with watery diarrhea (42). Allelic exchange with a counter-selectable suicide vector harboring *sacB* (pRE112; Cm^R^; (111)) was used to generate in-frame deletions of *invA_1a_* and *invA_1b_*. Deletion cassettes were amplified from *P. alcalifaciens* 205/92 genomic DNA by overlap extension PCR using oligonucleotides listed in Table S2 and ligated into pRE112. Resulting plasmids were electroporated into *E. coli* SY327λpir for sequence confirmation, then transferred by electroporation into *E. coli* SM10λpir (Kan^R^), followed by conjugation into *P. alcalifaciens* 205/92 WT (Tet^R^). Selection of conjugants was on LB agar containing tetracycline (10 µg/ml) and chloramphenicol (30 µg/ml). Resulting meridiploids were counter-selected by incubating overnight on LB agar containing 1% (w/v) tryptone, 0.5% (w/v) yeast extract and 5% (w/v) sucrose at 30°C. Sucrose-resistant clones were screened by PCR using gene-specific primers and primers flanking the recombination region to verify gene deletions. For *in trans* complementation of μ*invA*_1a_, the promoter region of *invF*_1a_ was fused to the open reading frame of *invA*_1a_ by overlap extension PCR using the oligonucleotides listed in Table S2 with *P. alcalifaciens* genomic DNA. The resulting amplicon was then ligated into the *Bam*HI/*Xma*I sites of pGEN-MCS (112) to generate pGEN-*invA*_1a_ which was then electroporated into *P. alcalifaciens* μ*invA*_1a_ bacteria. For fluorescence detection of *P. alcalifaciens*, bacteria were electroporated with pGEN-DsRed (113), which encodes for the red fluorescent protein variant, DsRed.T3, under the control of the *em7* promoter.

To generate transcriptional reporters for genes associated with T3SS_1a_, T3SS_1b_, and flagella, predicted promoter regions of putative regulators or structural components were amplified with oligonucleotides listed in Table S2 and cloned into pFU35 (114) upstream of the *luxCDABE* operon. Sequence-confirmed reporter plasmids were electroporated into *P. alcalifaciens* WT bacteria*. S*. Typhimurium SL1344 was the wild-type strain used in this study (115). The *S*. Typhimurium SL1344 translocator swap mutants, Δ*sipC*::*PasipC*_1a_ and *sipC*::*cipC*, have been described previously (50).

### Bacterial genome sequencing and assembly

A log phase culture of *P. alcalifaciens* 205/92 was centrifuged and the bacterial pellet resuspended in 1x DNA/RNA Shield (Zymo). Bacterial DNA was extracted and sequenced at Plasmidsaurus Inc. (Eugene, Oregon) using Oxford Nanopore Technologies (v14 library prep chemistry, R10.4.1 flow cell, basecalled using dna_r10.4.1_e8.2_5khz_400bps_sup@v4.2.0 model, primer- and adapter-trimmed) and Illumina (NextSeq 2000, 153 bp paired-end reads). Raw sequencing reads were deposited to ENA (BioProject PRJNA1100810). Raw Illumina reads were adapter and quality-trimmed using bbduk.sh v38.07, using options “ktrim=r k=23 mink=11 hdist=1 tpe tbo”. Nanopore reads were filtered using filtlong v0.2.1 against the trimmed Illumina reads, with options “--min_length 1000 --keep_percent 90 --target_bases 500000000 –trim --split 500”, selecting approximately 100x coverage of best Nanopore reads. All scripts used in the analysis can be found in https://github.com/apredeus/P_alcalifaciens.

Resulting filtered Nanopore reads were assembled using Trycycler v0.5.4, which uses manual curation of long-read assemblies to achieve a nearly perfect bacterial genome assembly. To this end, raw Nanopore reads were randomly sub-sampled to 50x depth and assembled 10 times each using the following long-read assemblers: flye v2.9.2-b1786, raven v1.8.1, and miniasm 0.3-r179. Following this, 25 assemblies (9 flye, 8 raven, and 8 miniasm) were selected for further analysis. Assembled contigs were clustered and evaluated; three supported clusters were retained, outlier contigs in each cluster were removed, and reconciled sequences generated. Additionally, Unicycler v0.5.0 was run in hybrid mode on both Nanopore and Illumina reads to recover small plasmids that could potentially be missed by long-read approaches because of size selection bias. This allowed the recovery of a small (∼4 kb) plasmid that was missed by long read-only assembly.

The final assembly consisted of 1 chromosome and 3 plasmids. The chromosome was rotated to match the start of RefSeq assembly GCF_002393505.1 (strain FDAARGOS_408, representative *P. alcalifaciens* genome). Following this, polishing with the trimmed Illumina reads using Polypolish v0.5.0 (Wick and Holt, 2022) was done to correct the remaining small indels and single-nucleotide polymorphisms. The resulting assembly was submitted to NCBI (GenBank Accession No. GCA_038449115.1). To annotate the genome, Bakta v1.9.1 was run with settings “--complete -- compliant --genus Providencia --species alcalifaciens --locus-tag PA205 --keep-contig-headers”. To annotate candidate prophage regions, PHASTER online service (https://phaster.ca/, accessed on March 15th, 2024) together with the manual curation using individual protein annotations from Bakta was used.

For the comparative genome analysis, all of the *P. alcalifaciens* genome assemblies available via Genbank on March 10th, 2024 were downloaded as finished assemblies; entries marked as anomalous by NCBI were excluded. Additionally, representative genomes of *P. rustigianii* (strain 52579_F01, assembly GCA_900637755.1) and *P. rettgeri* (strain AR_0082, assembly GCA_003204135.1) were downloaded to be used as potential outgroups. In total, 51 assemblies were downloaded; of these, 13 were marked as “Complete genome”, 28 were marked “Contig”, and another 8 were marked “Scaffold” (Table S1). Using snippy v4.6.0 in contig mode, and our assembly of strain 205/92 as a reference, all other assemblies were mapped and variant-called. Assemblies were evaluated for quality, and assemblies with more than 60% of the reference genome covered were removed (Table S1). Following this, snippy-core utility v4.6.0 was used to create core genome alignment of the remaining genomes.

Pairwise SNP distances were calculated using snp-dist v0.8.2 using the AGCT-only core genome alignment. Constant sites in the full core genome alignment were identified using snp-sites v2.5.1. In order to determine the plasmid coverage, *in silico* reads generated by Snippy from assembled contigs (single-end, 250 bp) were mapped to our assembly of *P. alcalifaciens* strain 205/92 using bwa v0.7.17-r1188, and samtools v1.18 “coverage” command was used to calculate the coverage of individual plasmids. Plasmids with 80% coverage were classified as fully present; plasmids with 40-80% coverage were classified as partially present.

IQTree v2.2.4 was run using constant sites defined above, and with the options “-redo -ntmax 16 -nt AUTO -st DNA -bb 1000 -alrt 1000”. Best-fit model was selected by Bayesian information criterion. The resulting tree, plasmid presence, and SNP distances were visualized using R 4.3.3 with packages ggtree (v3.10.1), treeio (v1.26.0), tidytree (v0.4.6), and pheatmap (v1.0.12).

BRIG v0.95 and NCBI blast 2.7.1+ were used to align and produce the circular visualization of the chromosome and the largest plasmid. To be used in BRIG, all of the multi-fasta genome assemblies were converted to a single fasta using a custom script. All scripts used for the analysis and visualization are available at https://github.com/apredeus/P_alcalifaciens.

### Bacterial growth curves

*P. alcalifaciens* was grown overnight in LB-Miller broth (Difco) for 16-18 h with shaking (220 rpm) at 37°C. Cultures were back-diluted into 10 mL LB-Miller broth in a 125 ml Erlenmeyer flask to a starting optical density at 600 nm (OD_600_) of 0.1 and grown with aeration (220 rpm) at 37°C. Bacterial growth curves were generating by sampling 1 mL of subculture in cuvettes and measuring the (OD_600_) in a BioRad SpartSpec Plus spectrophotometer every hour.

### Bacterial luminescence

Bacteria were subcultured as described above for 12 h. Each hour, 150 µl of subculture was transferred, in duplicate, to a white flat-bottom 96-well polystyrene microplate (Corning Costar) sealed with polyester Axyseal film (Axygen Inc.). Luminescence was measured using a Tecan Infinite M1000 plate reader.

For quantification of bacterial luminescence upon infection of mammalian and insect cells, cells were seeded in 6-well plates: (1) HCT116 cells at 4×10^5^ cells/well on rat tail collagen ∼40-44 h prior to infection; (2) S2 cells at 2×10^6^ cells/well ∼24 h prior to infection. Infections were as described above with *P. alcalifaciens* subcultures. At the required timepoint, infected monolayers were washed twice with HBSS, collected in 1 ml sterile double distilled water using a cell scraper, transferred to a 1.5 ml Eppendorf tube, vortexed, then centrifuged at 8,000xg for 2 min to pellet bacteria. The supernatant was carefully removed and discarded, the pellet resuspended in 100 µl PBS and transferred to a white flat-bottom 96-well polystyrene microplate (Corning Costar). Luminescence was measured using a Tecan Infinite M1000 plate reader.

### Motility assays

Overnight cultures of *P. alcalifaciens* WT*, S.* Typhimurium SL1344 WT and Δ*flgB* mutant (Ibarra et al., 2010) were inoculated using a sterile pipette tip into the center of Petri dishes containing LB plus 0.3% (w/v) agar using a sterile pipette, piercing approximately half-way through the semi-solid agar. Plates were incubated overnight at 37°C.

### Secretion assays and mass spectrometry

Two 10 mL subcultures were grown for 4 h with shaking for each strain, as described above, and pelleted for 15 min at 16,000x *g*. Supernatants were pooled and filtered with a 0.22 µm low-protein binding Acrodisc filter (Whatman). Proteins were precipitated in 10% (w/v) trichloroacetic overnight at 4°C. Protein precipitates were collected by centrifugation at 16,000x *g* for 15 min at 4°C and pellets were washed with cold acetone, dried, and resuspended in 200 µl 1.5X SDS-PAGE sample buffer. Secreted proteins were separated on a 10% SDS-PAGE gel and visualized by Coomassie Brilliant Blue stain (Fisher). For protein identification, secreted proteins were separated on 4-15% gradient SDS-PAGE gels (BioRad), stained with GelCode Blue (Thermo) and bands of interest were excised and sent to Stanford University Mass Spectrometry (SUMS) for protein identification by LC/MS/MS. Following in-gel tryptic digestion reconstituted samples were analyzed using a nanoAcquity UPLC (Waters) coupled to an Orbitrap Q-Exactive HF-X (RRID:SCR_018703) mass spectrometer.

### Tissue culture

HCT116 cells (human colorectal carcinoma epithelial), HeLa (human cervical carcinoma), THP-1 (human monocytes) and J774A.1 (mouse macrophage-like) cells were purchased from ATCC and used within 15 passages of receipt. HCT116 cells were maintained in McCoy’s medium 5A (Iwakata and Grace Modification, Corning) supplemented with 10% (v/v) heat-inactivated fetal calf serum (FCS, Invitrogen). HCT116 *CASP1*^-/-^ and *CASP4*^-/-^ knockout (KO) cells were generated using CRISPR-Cas9 technology by Synthego (www.synthego.com). *CASP1*^-/-^ KO clones G15 and I17, and *CASP4*^-/-^ KO clones J3 and N8 were derived by single cell expansion. Premature termination of the respective genes was verified by DNA sequencing. Where indicated, HCT116 cells were treated with IFNy (PeproTech) at 50 ng/ml for 16-18 h. HeLa cells were maintained in Eagle’s minimal essential medium (Corning) containing 2 mM L-glutamine, 1 mM sodium pyruvate and 10% (v/v) heat-inactivated FCS. THP-1 cells were maintained in RPMI1640 medium (Corning) containing 1 mM sodium pyruvate, 2 mM L-glutamine, 10 mM Hepes and 10% (v/v) heat-inactivated FCS. J774A.1 cells were maintained in Dulbecco’s modified Eagle’s medium (4.5 g/L glucose, Corning) containing 4 mM L-glutamine and 10% (v/v) heat-inactivated FCS. All mammalian cell lines were maintained at 37°C in 5% CO_2_. *Drosophila* S2 cells were purchased from Invitrogen and used up to passage number 20. Cells were maintained in Schneiders’ medium (Invitrogen) with penicillin-streptomycin and 10% (v/v) heat-inactivated FCS. S2 cells were maintained at 26°C without CO_2_. Cells were seeded in 24-well tissue culture treated plates (Nunc) at the following densities: (i) HCT116, 1×10^5^ cells/well ∼44 h prior to infection on rat tail collagen (Corning), (ii) HeLa, 5×10^4^ cells ∼24 h prior to infection, (iii) THP-1, 2.5×10^5^ cells/well ∼48 h prior to infection in the presence of 200 nM phorbol 12-myristate 13-acetate (PMA, LC Laboratories), growth media was replaced with PMA-free media 4 h prior to infection, (iv) J774A.1, 2×10^5^ cells/well ∼ 24 h prior to infection, (iv) S2, 5×10^5^ cells/well ∼24 h prior to infection, growth media was changed to penicillin-streptomycin-free media 4 h prior to infection.

### Gentamicin protection and CHQ resistance assays

To enumerate intracellular bacteria, cells were seeded and infected as described above, and subject to a gentamicin protection assay as described (66). *P. alcalifaciens* strains were subcultured for 3.5-4 h as detailed above. One mL of subculture was pelleted at 8,000x *g* for 90 sec, resuspended in 1 mL Hanks’ balanced salt solution (HBSS, Corning), and diluted 1:100 in HCT116 growth media or 1:1000 in THP-1 or S2 growth media. Because *P. alcalifaciens* are poorly motile (Figure S2), we centrifuged late log-phase cultures onto host cells to promote bacteria-host cell association, a technique typically used for non-motile or poorly motile bacteria (59, 61, 65). Without this centrifugation step, no *P. alcalifaciens* were internalized into non-phagocytic (HCT116) cells after 30 min of co-incubation (our unpublished results). One mL of diluted culture was centrifuged onto cells at 500x *g* for 5 min at room temperature (t_0_) (multiplicity of infection (MOI) of ∼150 for HCT116, ∼ 10 for THP-1, ∼ 5 for S2), then monolayers were incubated for a further 25 min at 37°C (HCT116, THP-1) or 26°C (S2). Extracellular bacteria were removed at 30 min p.i. by washing 3 times with 1 mL HBSS, then incubating in growth media containing 100 µg/mL gentamicin at 37°C (HCT116, THP-1) or 26°C (S2). At 1 h p.i., monolayers were washed once with 1X PBS and solubilized in 0.2% (w/v) sodium deoxycholate (NaDOC). Internalized bacteria and subculture inocula were serially diluted and plated on LB agar for CFU enumeration. Invasion efficiency was quantified as number of internalized bacteria/inoculum x 100%. To measure intracellular replication over time, the multiplicity of infection (MOI) for the Δ*invA*_1a_ mutant was increased by 2-3-fold for HCT116 infections (MOI of 300-600) so that an approximately equivalent number of WT and Δ*invA*_1a_ bacteria were internalized at 1 h p.i. Increasing the MOI of Δ*invA*_1a_ bacteria up to 10-fold did not compensate for the invasion defect of a Δ*invA*_1a_ mutant in S2 cells, however. Media containing 100 µg/ml gentamicin was added from 30-90 min p.i., then replaced with media containing 10 µg/ml gentamicin for the remaining time. Monolayers were solubilized at 4 h, 8 h, and 12 h p.i., serially diluted and plated as described above. To determine the proportion of intracellular bacteria in the cytosol of eukaryotic cells, the chloroquine (CHQ) resistance assay was used as previously published (65, 66). The concentration of CHQ was 400 µM for all cell types.

### Inside/outside microscopy assay

Extracellular and intracellular bacteria were distinguished by staining with anti-*Providencia* antibodies in the absence of any permeabilizing agents. Cells were seeded on acid-washed 12 mm diameter glass coverslips (#1.5 thickness, Fisherbrand) in 24-well plates and infected with *P. alcalifaciens* WT, Δ*invA*_1a_ or Δ*invA*_1b_ bacteria carrying pGEN-DsRed.T3. At 1 h and 8 h p.i., monolayers were fixed in 2.5% paraformaldehyde at 37°C for 10 min, then washed thrice in PBS. Extracellular bacteria were labelled with rabbit polyclonal anti-*P. alcalifaciens* antibody (kindly provided by Dr M. John Albert) diluted 1:250 in PBS containing 10% (v/v) normal goat serum (Invitrogen) for 15 min at room temperature. Monolayers were washed thrice in PBS, once in PBS containing 10% (v/v) normal goat serum, then incubated with goat anti-rabbit Alexa-Fluor 488 secondary antibodies diluted 1:300 in PBS containing 10% (v/v) normal goat serum at room temperature for 15 min (150 µl per well). After three washes in PBS, host cell nuclei were labelled with Hoechst 33342 for 1 min (Invitrogen, 1:10,000 dilution in water) and coverslips mounted in Mowiol (Calbiochem) on glass slides. Samples were viewed on a Leica DM4000 upright fluorescence microscope. A bacterium was scored as extracellular if it fluoresced red and green, or intracellular if it fluoresced red only.

### Electron microscopy

SEM and TEM sample preparation and imaging was performed at the Franceschi Microscopy and Imaging Center at Washington State University using standard techniques. For SEM, HeLa and HCT116 cells were seeded on Thermanox plastic coverslips (Nunc), infected with *P. alcalifaciens* as described above and at 20 min p.i., washed once in HBSS and fixed in 2% paraformaldehyde/2% glutaraldehyde in 0.1 M cacodylate buffer pH 7.2. overnight at 4°C. Glutaraldehyde-fixed cells were dehydrated in an ethanol series and dried at the critical point in CO_2_. The samples were sputter coated with platinum/palladium to 2.5 nm thickness and examined on a FEI Quanta 200F scanning electron microscope.

For TEM, HeLa cells were seeded in T-25 tissue culture flasks (Nunc) at 7 ×10^5^ cells per flask the day before infection; HCT116 cells were seeded on rat tail collagen at 4×10^5^ cells/well in 6-well plates ∼40 h prior to infection; THP-1 cells were seeded in the presence of PMA at 1×10^6^ cells/well in 6-well plates 2 days prior to infection; S2 cells were seeded at 2×10^6^ cells/well in 6-well plates. Monolayers were infected with *P. alcalifaciens* as described above and at 60-90 min p.i. gently washed thrice with PBS and treated with 0.25% trypsin (Corning) to detach HCT116 and HeLa cells, or TrypLE Select (Gibco) to dislodge THP-1 cells. S2 cells were dislodged with a cell scraper (Sarstedt). Cells were collected, centrifuged at 400xg for 5 min, the supernatant discarded, and the cell pellet gently resuspended in fixative. Processing and imaging were as described previously (Du et al., 2016).

### Cytotoxicity assays

Prior to infection, medium was replaced on HCT116 cells and THP-1 cells to phenol-red free RPMI1640 (Corning) containing 10% (v/v) heat-inactivated FCS. Infections and subsequent steps were in phenol-red free media. Cells were infected as described above and supernatants collected at the indicated times, then centrifuged at 500 xg for 5 min to pellet cellular debris. Cell-free supernatants were collected and stored at −80°C until analysis. Lactate dehydrogenase released into the supernatant, a measure of the loss of plasma membrane integrity, was quantified using the Cytotox96 Assay Kit (Promega) according to the manufacturer’s instructions.

### Cytokine quantification by ELISA

HCT116 culture supernatants were collected at the indicated times, centrifuged at 500xg for 5 min, then the cell-free supernatants were stored at −80°C until analysis. IL-18 was quantified by sandwich ELISA as previously described (67). Mouse anti-human IL-18 monoclonal (125-2H) and rat anti-human IL-18 monoclonal (159-12B) biotin were purchased from MBL. Human IL-8 was quantified by DuoSet ELISA (R&D Systems) as per the manufacturer’s instructions.

### Immunoblotting

For analysis of whole cell lysates, adherent cells were washed once in PBS prior to lysis in boiling 1.5x SDS-PAGE sample buffer. Proteins were separated by SDS-PAGE and transferred to 0.45 µm pore-size nitrocellulose membranes (GE Healthcare Life Sciences). Membranes were blocked at room temperature for 1 h with Tris-buffered saline (TBS) containing 5% (w/v) skim milk powder and 0.1% (v/v) Tween-20 (TBST-milk), then incubated with the following primary antibodies overnight at 4°C: rabbit polyclonal anti-caspase-1 (A-19) (1:2,000; Santa Cruz Biotechnology), mouse monoclonal anti-caspase-4 (4B9) (1:2,000; MBL), mouse monoclonal anti-β-actin (8H10D10) (1:20,000; Cell Signaling Technology). Blots were then incubated with anti-rabbit IgG or anti-mouse IgG horseradish peroxidase (HRP)-conjugated secondary antibodies (1:10,000; Cell Signaling) in TBST-milk for 1-2 h at room temperature, followed by Supersignal West Femto Max Sensitivity ECL Substrate (Thermo). Chemiluminescence was detected using a GE Healthcare AI600 imager.

### Quantitative real-time PCR (qPCR)

To evaluate the effect of IFNγ priming on the expression of *CASP1*, *CASP4*, *CASP5*, *IL18*, *IL1B* in HCT116 cells we used Luminaris Color HiGreen qPCR Master Mix (Thermo Scientific) and a C1000 Touch Thermal Cycler, CFX96 Real-Time System (Bio Rad) with validated oligonucleotide primer pairs as we have described previously (67). Relative gene expression levels were quantified based on quantification cycle (Cq) values and normalized to the reference gene, ribosomal phosphoprotein P0 (*RPLP0*). The expression of each gene was calculated using the 2^-ΔCq^ method.

### D. melanogaster infections

*P. alcalifaciens* were grown overnight at 37°C for 16-18 h with shaking (220 rpm) in LB-Miller broth. Cultures (1 ml) were centrifuged at 8,000xg for 90 sec, the bacterial pellet washed twice in an equal volume of sterile PBS and diluted 1,000-3,000-fold in sterile PBS. Two-to-seven-day old *Wolbachia*-free adult male *w^1118^* flies (Bloomington Drosophila Stock Center #5905) were anesthetized with CO_2_ and injected with 23 nl of diluted culture (∼10-30 CFU/fly) or PBS vehicle control. Flies were maintained on standard cornmeal food at 25°C and 65% relative humidity and surviving flies were counted every 2-6 h p.i. In each experiment, ∼50 flies were injected for each condition and survival studies were repeated at least twice for each infectious dose (∼10 and ∼30 CFU/fly).

### Bovine ligated jejuno-ileal loop infections

Jersey, Holstein, or cross-bred calves were obtained from North Carolina State University or University of Wisconsin farm herds. Calves were separated from the dam and transferred to AAALAC-approved large animal housing facilities by 1 day of age. Calves were administered either colostrum or colostrum replacer and adequate passive transfer was estimated by measurement of serum total protein. Calves were treated for 3 days with ceftiofur (4-6 mg/kg SC q24h) and/or flunixin meglumine (50 mg/kg IV q24h) if needed based on clinical condition upon arrival. Calves were fed milk replacer at 10-20% body weight per day with free choice access to water and hay.

At 3 to 6 weeks of age, calves were anesthetized with intravenous propofol and maintained on isoflurane inhalant for ligated jejuno-ileal loop surgery as previously described (116). Briefly, calves were placed in left lateral recumbency, and a right flank incision was made. Sixteen to thirty-eight 4- to 6-cm loops were tied within the ileum and terminal jejunum leaving 1-cm spacers between adjacent loops. Prior to inoculation, loop lengths were recorded. Loops injected with vehicle only served as negative controls. Loops were infected individually with 3 ml LB (12 h incubation) or 2 ml PBS (2 h and 8 h incubation) containing approximately 10^9^ CFU of *P. alcalifaciens*. The intestine was returned to the abdomen, the incision was closed, and the calves were monitored under inhalant anesthesia for the duration of the experiment. At 2 h, 8 h, or 12 h p.i., the incision was opened, and each loop was individually excised. Calves were euthanized by intravenous pentobarbital.

In preparation for ligated loop infections, bacteria were grown shaking (225 rpm) overnight at 37°C in LB-Miller broth. For the 12 h infections, overnight cultures were subcultured 1:100 into LB-Miller broth and further incubated for 3.5-4 h at 37°C with shaking (225 rpm). Bacteria were washed twice and resuspended in PBS (2 h or 8 h) or LB (12 h) with bacterial normalization based on optical density (OD_600_) for a final inoculation dose of ∼10^9^ *P. alcalifaciens* per loop. Actual inoculum dose was determined by serial dilution and plating. Following loop excision, intestinal fluid and tissue samples were harvested and processed separately. Fluid volume was calculated by excising individual loops and weighing escaped luminal fluid on a sterile petri dish. Luminal fluid was then transferred to 1 mL sterile PBS to allow for bacterial enumeration. Intestinal tissue was washed twice in sterile PBS to remove non-adherent bacteria and ingesta and then added to 5 mL sterile PBS. For 2 h and 8 h infections, washed tissues were cut in half with one segment treated with gentamicin (50 µg/mL) for 30 minutes at 37°C to quantify intracellular bacteria, and the remaining half processed to quantify tissue-associated bacteria. After gentamicin treatment, tissues were washed twice with PBS to remove remaining antibiotics. Samples were subsequently homogenized, serially diluted in PBS, and plated for CFU enumeration.

The Institutional Animal Care and Use Committees of North Carolina State University and University of Wisconsin-Madison approved all animal experiments (NCSU protocol numbers 15-047-B and 17-186-B; UW-Madison protocol number V006249). All animal experiments were performed in accordance with the PHS “Guide for the Care and Use of Laboratory Animals” in AAALAC-approved animal facilities.

### Histopathology

Intestinal samples were fixed in neutral 10% buffered formalin, processed for paraffin embedding, sectioned (5 μm), and stained with hematoxylin and eosin for histologic analyses. Tissues were assessed and scored by an American College of Veterinary Pathology (ACVP) board-certified pathologist with a scoring system derived from previously published rubrics (117, 118){Citation}. The following criteria were scored from 0 to 4: lamina propria neutrophil accumulation, submucosal edema, epithelial damage, crypt abscess, villus blunting, cell death.

| Score | Lamina propria neutrophil accumulation | Submucosal edema | Epithelial damage | Crypt abscess | Villus blunting | Cell death |
| --- | --- | --- | --- | --- | --- | --- |
| 0 | 0-5/hpf | No change (0-5) | No change | None | Normal villi | Absence of cells with morphological features of death |
| 1 | 6-20/hpf | 5-10 | Detectable (<10%) | Detectable (<10%) | Few villi mildly blunted | Few sporadic necrotic/apoptotic cells detected |
| 2 | 21-60/hpf | 11-20 | Mild (10-20%) | Mild (10-20%) | Most villi mildly blunted | Minimal number necrotic/apoptotic cells detected |
| 3 | 61-100/hpf | 21-40 | Moderate (21-40%) | Moderate (21-40%) | Most villi moderately blunted | Moderate number necrotic/apoptotic cells detected |
| 4 | >100/hpf | >40 | Marked (>40%) | Marked (>40%) | All villi severely blunted | Large number necrotic/apoptotic cells detected |

### Statistical analysis

Statistical analysis was performed using GraphPad Prism 6 software. Statistical significance of comparisons between treatment groups was determined using either an unpaired, two-tailed Student’s *t* test, or for group analysis, using one-way analysis of variance (ANOVA) followed by Dunnett’s multiple-comparison test.

## Acknowledgements

We thank Dr Allis Chien and Dr Ryan Leib at Stanford University Mass Spectrometry (SUMS) facility for mass spectrometry analysis; the staff at the Franceschi Microscopy and Imaging Center (FMIC) at WSU for their expertise and use of electron microscope facilities; Patricia Reis for her technical assistance in the initial stages of this project; Dr J. Michael Janda for his help in sourcing the *P. alcalifaciens* 205/92 strain; Dr M. John Albert for providing anti-*P. alcalifaciens* antibodies; Dr Harry Mobley and Stephanie Himpsl for providing pGEN-DsRed.T3 and pGEN-MCS plasmids; and Dr Petra Dersch for providing the pFU35 plasmid.

Research reported in this publication was supported, in part, by a Burroughs Wellcome Fund Investigators in the Pathogenesis of Infectious Diseases Award to L.A.K., and the National Institute of Allergy and Infectious Diseases (NIAID) of the National Institutes of Health (NIH) under award numbers R21AI130645 to L.A.K. and R01AI134766 to L.A.K and B.A.V. The funders had no role in study design, data collection and interpretation, or the decision to submit the work for publication. Jessica A. Klein is currently employed by Genentech Inc. Genentech Inc. was not involved in this study or its design.

## Supplementary information

**Table S1.** *P. alcalifaciens* genome information used for analysis. Table S2. Oligos used for cloning.

**Figure S1. Alignment of p128kb.** BRIG diagram of p128kb alignment from the ten *P. alcalifaciens* strains that harbor p128kb and are not shown in Figure 1B (PAL-3, F90-2004, wls1935, JBDGAAB-19-0025, 2019-04-27799-1-2, 2019-04-29034-1-3, 2019-04-28369-1-2, 2019-01-3283-1-1, 2019-04-28370-1-5, GCA_958348825.1). p128kb of *P. alcalifaciens* 205/92 presented in this work is used as a reference. Exact coordinates of T3SS_1a_ are provided in Table S1.

**Figure S2. *P. alcalifaciens* is poorly motile.** (A) Schematic of the *P. alcalifaciens* 205/92 genomic region containing flagella-associated genes. (B) Promoter activity of flagella regulators and structural components reported as 10^4^ relative luminescence units (RLUs). Data are from ≥3 independent experiments (mean ± SD). (C) Swimming motility of *P. alcalifaciens* compared to *S.* Typhimurium WT and a non-motile *S.* Typhimurium Δ*flgB* mutant. Overnight bacterial cultures were inoculated into semi-solid LB agar plates and incubated overnight at 37°C.

