## Supplementary figures and images for "Pathogenic diversification of the gut commensal *Providencia alcalifaciens* via acquisition of a second type III secretion system"

### Supplemental Figures

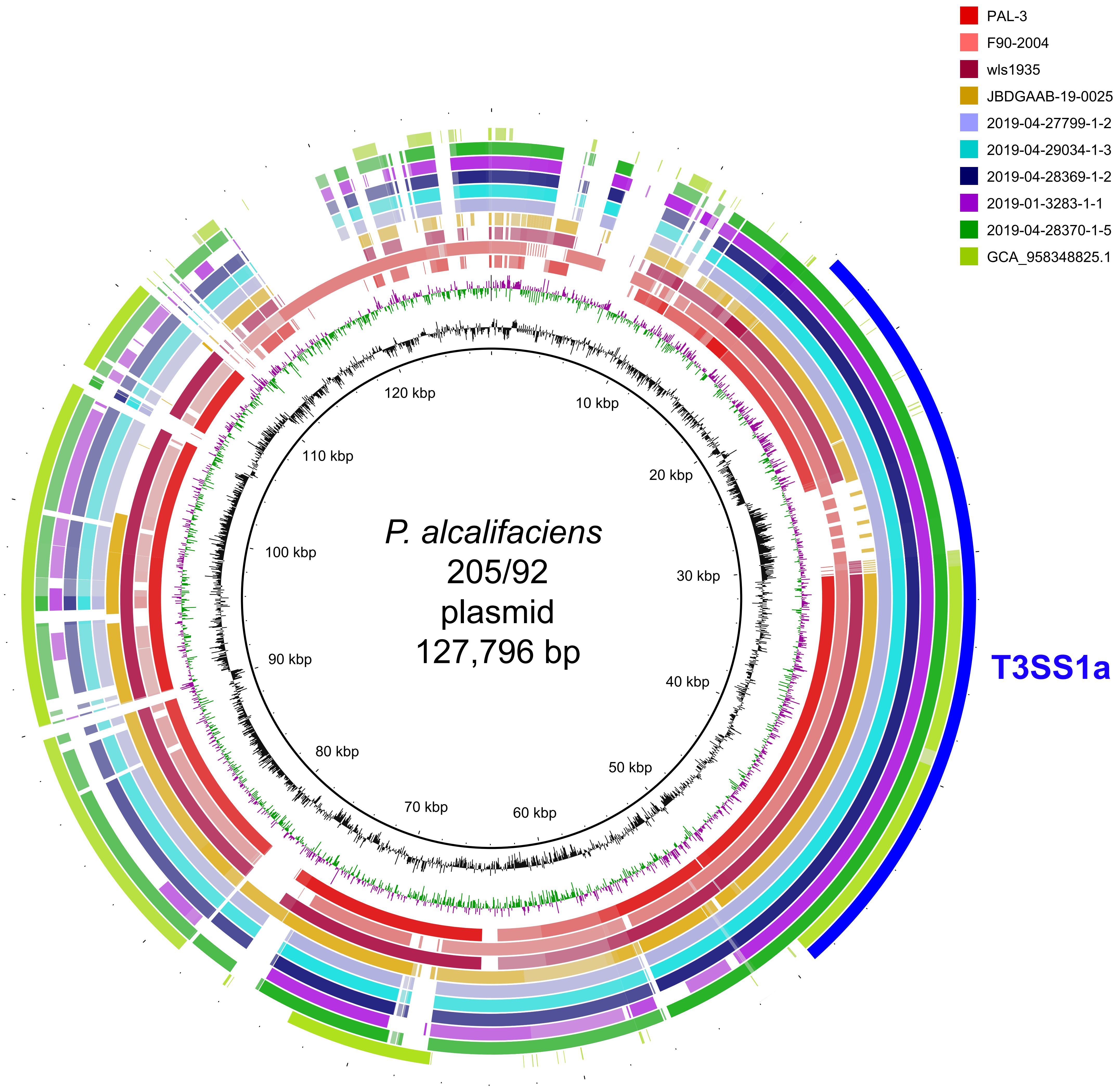

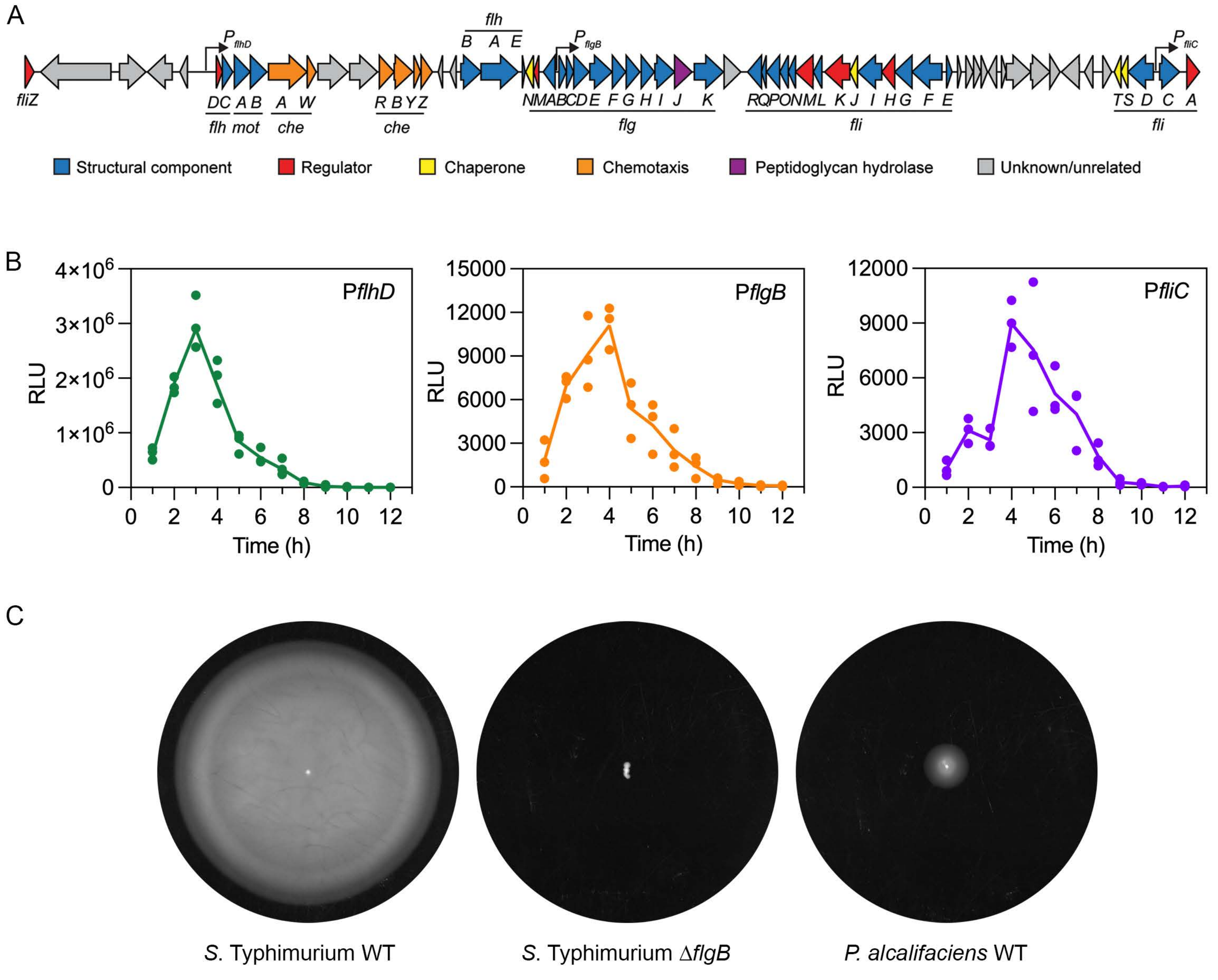
